# Intranasal Human NSC-Derived EVs Therapy Can Restrain Inflammatory Microglial Transcriptome, and NLRP3 and cGAS-STING Signaling, in Aged Hippocampus

**DOI:** 10.1101/2025.03.02.641082

**Authors:** Leelavathi N. Madhu, Maheedhar Kodali, Shama Rao, Sahithi Attaluri, Raghavendra Upadhya, Goutham Shankar, Bing Shuai, Yogish Somayaji, Shruthi V. Ganesh, Vignesh S. Kumar, Jeswin E. James, Padmashri A. Shetty, Avery LeMaire, Xiaolan Rao, James J Cai, Ashok K. Shetty

## Abstract

Neuroinflammaging, a moderate, chronic, and sterile inflammation in the hippocampus, contributes to age-related cognitive decline. Neuroinflammaging comprises the activation of the nucleotide-binding domain, leucine-rich repeat family, and pyrin domain-containing 3 (NLRP3) inflammasomes, and the cyclic GMP-AMP synthase (cGAS)-stimulator of interferon genes (STING) pathway that triggers type 1 interferon (IFN-1) signaling. Studies have shown that extracellular vesicles from human induced pluripotent stem cell-derived neural stem cells (hiPSC-NSC-EVs) contain therapeutic miRNAs that can alleviate neuroinflammation. Therefore, this study examined the effects of late middle-aged (18-month-old) male and female C57BL6/J mice receiving two intranasal doses of hiPSC-NSC-EVs on neuroinflammaging in the hippocampus at 20.5 months of age. Compared with animals receiving vehicle treatment, the hippocampus of animals receiving hiPSC-NSC-EVs exhibited reductions in astrocyte hypertrophy, microglial clusters, and oxidative stress, along with elevated expression of antioxidant proteins and genes that maintain mitochondrial respiratory chain integrity.

Moreover, hiPSC-NSC-EVs therapy decreased the levels of various proteins involved in the activation of the NLRP3 inflammasome, p38/mitogen-activated protein kinase, cGAS-STING-IFN-1, and Janus kinase and signal transducer and activator of transcription signaling pathways. Furthermore, in vitro assays using genetically engineered RAW cells and hiPSC-NSC-EVs, with or without targeted depletion of specific miRNAs, demonstrated that miRNA-30e-3p and miRNA-181a-5p, both present in hiPSC-NSC-EVs, can significantly inhibit the activation of the NLRP3 inflammasome and the STING pathway, respectively. Additionally, single-cell RNA sequencing conducted 7 days post-treatment revealed that hiPSC-NSC-EVs induce widespread transcriptomic changes in microglia, including increased expression of numerous genes that enhance oxidative phosphorylation and reduced expression of abundant genes that drive multiple proinflammatory signaling pathways. These changes mediated by hiPSC-NSC-EVs were also associated with improved cognitive and memory function. Thus, intranasal hiPSC-NSC-EVs therapy in late middle age can effectively diminish proinflammatory microglial transcriptome and signaling cascades that drive neuroinflammaging in the hippocampus, contributing to better brain function in old age.

## 1. INTRODUCTION

Improved human life expectancy over the past century has brought about longer lives and a surge in age-related ailments. Particularly, neurodegenerative diseases, including Alzheimer’s disease (AD), have emerged as the leading causes of morbidity worldwide [1–3]. Hence, effective strategies are urgently needed to promote successful aging, characterized by older adults maintaining better cognitive function, staying engaged in social activities, and displaying minimal signs of age-related diseases [4–6].

Aging is associated with reduced ability to adapt to new stimuli and cognitive decline in a significant proportion of individuals [7]. Studies using animal models and human post-mortem brain tissues have suggested that cognitive impairments associated with aging are often connected to various adverse changes in the brain, particularly in the hippocampus [6, 8–9]. The age-related changes include increased oxidative stress, mitochondrial dysfunction, and neuroinflammaging. Neuroinflammaging, a sterile and moderate chronic neuroinflammation, increases substantially in individuals who develop neurodegenerative disorders. It is characterized by increased concentrations of reactive oxygen species (ROS), associated with mitochondrial dysfunction [10–11] and the activation of several neuroinflammatory signaling cascades. These include the activation of the nucleotide-binding domain leucine-rich repeat (NLR) family pyrin domain-containing 3 (NLRP3) inflammasomes [12–13] and the cyclic GMP-AMP synthase (cGAS) and the stimulator of interferon genes (STING) pathway, which leads to chronic type I Interferon (IFN) signaling in the aged brain [14].

NLRP3 is predominantly expressed in microglia, the resident immune cells in the brain [15].

Activation of NLRP3 inflammasomes by danger-associated molecular patterns (DAMPs, e.g., increased ROS) in microglia increases the secretion of proinflammatory cytokines, such as interleukin-1 beta (IL-1β) and IL-18. These cytokines can induce pyroptosis in neighboring neural cells and activate the downstream p38 mitogen-activated protein kinase (p38/MAPK) signaling pathway via myeloid differentiation primary response 88 (Myd88) and the small GTPase rat sarcoma virus (Ras) [16–17]. This process can contribute to a chronic state of neuroinflammation with the continuous release of higher concentrations of multiple proinflammatory cytokines. On the other hand, activation of the cGAS-STING pathway in microglia, typically triggered by double-stranded DNA (dsDNA) released from damaged cells, leads to upregulation of type-I IFN production [14, 17], leading to the activation of Janus kinase and signal transducer and activator of transcription (JAK-STAT) signaling culminating in transcription of numerous interferon-stimulated genes (ISGs) [14, 18]. Thus, it is apparent that increased oxidative stress, mitochondrial dysfunction, and microglia-mediated chronic neuroinflammation in the aged brain increase the susceptibility to develop conditions such as mild cognitive impairment (MCI) or AD [5, 6, 19]. Therefore, strategies that restrain detrimental microglia-mediated inflammatory signaling cascades in the aging brain are needed to maintain better cognitive and mood function in old age. However, treatments capable of restraining such neuropathological changes associated with cognitive and memory impairments in late middle or old age have yet to be identified.

Previous studies have shown some beneficial effects of intracerebral neural stem cell (NSC) grafting in models of brain aging [20–21] and AD [22–23]. However, the benefits of NSC grafting are believed to be primarily due to bystander effects (i.e., paracrine actions of their secretome), which include antiinflammatory, neuroprotective, and neuroreparative activity [24–25]. From this perspective, extracellular vesicles (EVs) from human induced pluripotent stem cell (hiPSC)-derived NSCs, which retain most of the therapeutic effects of parental cells, have attracted attention for developing cell-free therapies. Also, intranasally (IN) administered hiPSC-NSC-EVs efficiently target microglia and astrocytes throughout the brain [26–27] and have been shown to induce beneficial transcriptomic changes in conditions such as AD [17]. Furthermore, the cargo (miRNAs and proteins) carried by these EVs has been previously validated for its ability to mediate antioxidant, antiinflammatory, and neuroprotective effects using *in vitro* and *in vivo* models [17, 26, 28–30]. Therefore, using a mouse model, this study investigated whether IN administration of hiPSC-NSC-EVs in late middle age can significantly reduce oxidative stress and curb microglia-mediated neuroinflammaging in the hippocampus. The results provide new evidence that hiPSC-NSC-EVs therapy in late middle age can diminish the proinflammatory transcriptome of microglia, leading to reductions in oxidative stress, mitochondrial dysfunction, and neuroinflammaging associated with better cognitive and memory function in old age. Robust antiinflammatory effects of hiPSC-NSC-EVs were apparent from diminished activation of signaling cascades such as the NLRP3, p38/MAPK, cGAS-STING, JAK-STAT, and IFN-1 pathways in the aged hippocampus.

## 2. MATERIALS AND METHODS

### 2.1. hiPSC-derived NSC cultures, purification and characterization of hiPSC-NSC-EVs

The procedures for generating NSCs from hiPSCs (Wisconsin International Stem Cell Bank (IMR90-4), passaging of hiPSC-NSCs, isolation of EVs from passage 11 (P11) hiPSC-NSCs, and characterization of the number, size, and markers of hiPSC-NSC-EVs are detailed in our previous reports [17, 26, 28] and the supplemental file.

### 2.2. Animals and Study Design

The study comprised two cohorts of C57BL/6 mice: young adult (3 months old) and late middle-aged (18 months old). We chose 18 months old mice, as this mouse age is approximately equivalent to a 60-year-old human [31]. The mice were purchased from Jackson Laboratories (Bar Harbor, Maine, USA) and housed with ad libitum access to food (4% fat diet) and water (n=131; 61 males and 70 females).

The Institutional Animal Care and Use Committee of Texas A&M University approved all studies conducted in this investigation. Male and female late middle-aged mice were randomly assigned to either the vehicle group (aged-Veh; n=31 [13 males and 18 females]) or the EVs group (aged-EVs; n=30 [12 males and 18 females]) for long-term studies. For single-cell RNA sequencing (sc-RNA-seq) studies on microglia, additional late middle aged male mice were recruited to the aged-Veh and aged-EVs groups (n=1/group). Furthermore, additional groups of 3-month-old mice (n=28, 14 males and 14 females) and 18-month-old mice (n=32, 16 males and 16 females) underwent neurobehavioral testing to assess cognitive status at 18 months of age, the timepoint chosen for the hiPSC-NSC-EVs treatment intervention in the study. Additionally, late-middle-aged mice were used to evaluate the incorporation of IN-administered PKH-26-labeled hiPSC-NSC-EVs into neural cells across various brain regions (n=8; 4 males and 4 females). Figure 1 [G] details the study design, including the time points for the IN-administration of hiPSC-NSC-EVs, neurobehavioral tests, and euthanasia.

**Figure 1:**
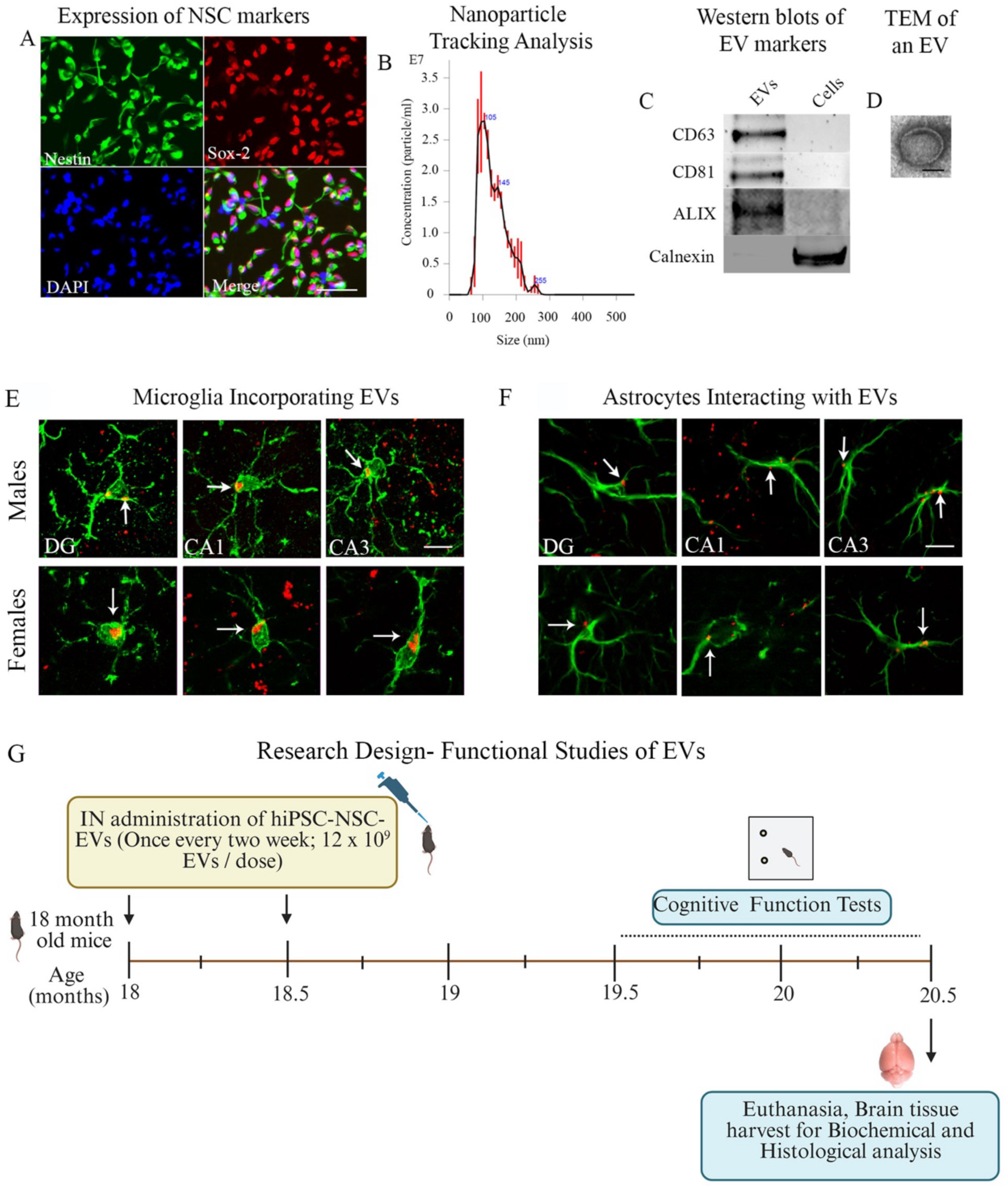
Characterization of extracellular vesicles (EVs) from human induced pluripotent stem cell-derived neural stem cells (hiPSC-NSCs), assessment of hiPSC-NSC-EVs biodistribution in the hippocampus and cerebral cortex of late middle-aged mice, and timeline of in vivo experiments. Image A illustrates that all cells in passage 11 NSCs derived from hiPSCs express NSC markers Nestin and Sox-2. Graph B shows the size and number of hiPSC-NSC-EVs measured with a NanoSight. The blots in C demonstrate EV-specific proteins CD63, CD81, and ALIX, and the absence of the deep cellular protein calnexin in hiPSC-NSC-EVs. The image in D shows the morphology and size of hiPSC-NSC-EVs as visualized by transmission electron microscopy. scale bars, A=100 µm, D=50 nm. Figure E illustrates the incorporation of EVs into IBA-1+ microglia in different hippocampal subregions (DG, CA1, and CA3) from male and female mice. Figure F illustrates the interaction of EVs with GFAP+ astrocytes in hippocampal subregions (DG, CA1, and CA3) from male and female mice. Scale bar, E-F =10 μm. Panel G illustrates the experimental design for long-term experiments showing the time points of hiPSC-NSC-EVs treatment, cognitive and mood function tests, euthanasia, and brain tissue harvest in late middle-aged male and female mice.

The animals in the aged-Veh or aged-EVs group for long-term studies received two doses of the vehicle or the hiPSC-NSC-EVs treatment (12 x 10^9^ EVs, ∼25 µg total protein), with each dose administered two weeks apart. The selection of dose was based on the results of our previous studies to reduce neuroinflammation in a lipopolysaccharide-induced peripheral inflammation model using young C57BL6 mice [29]. One month after the last dose, all animals underwent neurobehavioral testing. Following cognitive assessments, subgroups of animals were perfused with 4% paraformaldehyde, and collected brain tissues were processed for immunohistochemical studies (n=6-8/group) or euthanized to harvest fresh brain tissues for biochemical and molecular biological studies (n=6-8/group). The investigators conducting the behavioral tests and those performing immunohistochemical, immunofluorescence, biochemical, or molecular biological analyses were blinded to the group identities of animals or samples. Late middle-aged male and female mice recruited to biodistribution studies were euthanized 6 hours after receiving an IN administration of hiPSC-NSC-EVs labeled with PKH-26 (4 x 10^9^ EVs) to confirm the biodistribution of EVs into neural cells across various regions of the brain. Late middle-aged male mice recruited to sc-RNA-seq studies were euthanized 7 days after receiving a single IN dose of hiPSC-NSC-EVs (12 x 10^9^ EVs) to isolate live microglia.

### 2.3. Evaluation of the interaction of IN-administered hiPSC-NSC-EVs with microglia and astrocytes

The animals were perfused with 4% paraformaldehyde six hours after an IN administration of PKH-26-labeled hiPSC-NSC-EVs to visualize the interaction of IN-administered PKH-26-labeled hiPSC-NSC-EVs (fluorescing red) with neural cells. Thirty-micrometer-thick serial sections were cut using a cryostat, and every 15^th^ section through the entire forebrain from each animal (n=4/sex) was processed for ionized calcium-binding adaptor molecule 1 (IBA-1) or glial fibrillary acidic protein (GFAP) immunofluorescence to confirm the interaction or association of PKH-26-labeled hiPSC-NSC-EVs with microglia and astrocytes. The primary and secondary antibodies comprised anti-goat IBA-1 (1:1000, Abcam, Cambridge, UK), rabbit anti-GFAP (1:2000, Dako, Glostrup, Denmark), donkey anti-goat IgG conjugated to Alexa Fluor 488 (1:200, Invitrogen, Waltham, MA, USA), and donkey anti-rabbit IgG conjugated to Alexa Fluor 488 (1:200, Invitrogen). The supplemental file comprises the methods employed for IBA-1 and GFAP immunofluorescence.

### 2.4. Analyses of cognitive function

The study employed two neurobehavioral tests to measure cognitive function. A novel object recognition test (NORT) was used to investigate the competence for recognition memory, which relies on an interaction between the hippocampus and the perirhinal cortex [32]. Furthermore, an object location test (OLT) assessed the proficiency of animals in discerning subtle changes in their immediate environment, a cognitive function dependent on the integrity of the hippocampus [33]. The detailed protocols used for OFT, NORT, and OLT are available in our previous reports [16–17, 34–35] and in the supplementary file. In animals receiving vehicle or hiPSC-NSC-EVs treatment, these tests were performed between 19.5 and 20.5 months of age, along with age-matched naïve control animals.

### 2.5. Tissue processing, and immunohistochemical, immunofluorescence, and biochemical studies

After the completion of cognitive tests (i.e., at 20.5 months of age), the animals were euthanized, and the brain tissues were processed for immunohistochemical and biochemical analyses. The detailed methods employed for brain tissue collection, processing, immunohistochemistry, single/dual/triple immunofluorescence, and biochemical and molecular biological studies are available in our previous reports [16–17, 36–37] and the supplemental file.

### 2.6. Quantification of number and clusters of microglia in the hippocampus

The microglial clusters and microglial numbers in the hippocampus of male and female mice belonging to aged-Veh and aged-EVs groups (n=6/sex/group) were measured, using every 20^th^ section through the hippocampus. Microglial numbers were measured via stereological counting of IBA1+ cells using the optical fractionator method in the StereoInvestigator system (Microbrightfield, Williston, Vermont, USA). They were expressed as the number per 0.1 mm³ of hippocampal volume. The detailed stereological methods employed are described in our earlier reports [38–40]. We assessed the total number of microglial clusters for the hippocampus by counting every cluster (accumulation of 3 or more microglia) in serial sections (every 20^th^ through the entire hippocampus and then extrapolating the average number per section for the total number of sections through the hippocampus (n=6/group).

### 2.7. Measurement of oxidative stress markers, antioxidants, and nuclear factor erythroid 2-related factor 2 (NRF2)

Commercially available kits were employed to measure oxidative stress markers malondialdehyde (MDA) and protein carbonyls (PC) (Cayman Chemicals, Arbor, MI, USA), and NRF2 (Signosis, Santa Clara, CA, USA), in hippocampal lysates (n=6/sex/group). We followed the manufacturer’s instructions for performing these assays. Furthermore, to evaluate antioxidant status in the hippocampus, we measured superoxide dismutase (SOD) and catalase (CAT) concentrations in hippocampal lysates. The total protein concentration in the various tissue lysates was determined using a Pierce BCA Protein Assay Kit from ThermoFisher Scientific (Waltham, MA, USA). The concentrations of markers were normalized to the total protein content in their respective tissue lysates.

### 2.8. Measurement of genes linked to mitochondrial respiratory chain

We analyzed the expression of many genes regulating the mitochondrial respiratory chain in the hippocampus, using specific primers purchased from GeneCopoeia (Rockville, MD, USA) (n=6/sex/group). The measured genes encoding proteins relevant to the mitochondrial electron transport chain and oxidative phosphorylation include the *Ndufs6* and *Ndufs7* (Complex I), *Sdha* and *Sdhb* (Complex II), *Bcs1l* and *Cyc1* (Complex III); *Cox4i2* and *Cox7b* (Complex IV), *Atp6ap1* (Complex V) and *Slc25a10* (a gene encoding the mitochondrial dicarboxylate carrier protein). *Gapdh* was used as a housekeeping gene. The 2^delta-delta Ct (2^ddCT^) values for each gene were compared across different groups. All gene names mentioned in this manuscript have been expanded in the supplemental file (Suppl. Table 1).

### 2.9. Measurement of NLRP3 inflammasome genes, and NLRP3-ASC inflammasome complexes in microglia

We first performed qRT-PCR to measure the expression of genes *Nlrp3, Pycard, Casp1, Il1β,* and *Il18*. Then, to determine the extent of NLRP3-apoptosis-associated speck-like protein containing a caspase recruitment domain (ASC) inflammasome complexes and the percentages of microglia presenting NLRP3-ASC complexes, the brain tissue sections were processed for triple immunofluorescence to visualize IBA-1, NLRP3, and ASC in microglia. Using Z-section analysis in a Nikon confocal microscope or a Leica THUNDER 3D Imager, the number of NLRP3-ASC inflammasome complexes per unit area (∼216 μm^2^) and the percentages of IBA-1+ microglia co-expressing NLRP3-ASC inflammasome complexes were measured. For these quantifications, data collected from individual subfields of the hippocampus (DG, CA1, and CA3) were pooled across three sections/animal (n=6/sex/group) [16–17, 29]. The supplemental file comprises information on the antibodies employed and detailed methods.

### 2.10. Quantification of markers involved in the activation of NLRP3 inflammasome and p38/MAPK signaling cascades

The hippocampal lysates from male and female mice were processed to quantify the concentrations of various markers of NLRP3 inflammasome activation and p38/MAPK signaling (n=6/group), using commercially available individual enzyme-linked immunosorbent assay (ELISA) kits, as detailed in our previous studies [16–17, 41]. These include assay kits for 1) nuclear subunit of Nuclear factor kappa B (NF-kB p65; Aviva Systems Biology, San Diego, CA, USA), 2) NLRP3 (Abcam), 3) ASC (MyBioSource, San Diego, CA, USA), 4) cleaved caspase-1 (Abcam), 5) IL-18 (R&D Systems, Minneapolis, MN, USA), 6) IL-1β (R&D Systems, 7) MyD88 (Aviva Systems Biology), 8) Ras, (MyBioSource), 9) p38 MAPK (Cell Signaling, Danvers, MA, USA), 10) activator protein-1 (AP-1, Novus Biologicals, Centennial, CO; USA), and 11) tumor necrosis factor-alpha (TNF-α, R&D systems), and 12) IL-8 (Biomatik, Wilmington, DE, USA). The concentrations of individual proteins were normalized to 1 mg of total protein in hippocampal lysates.

### 2.11. Quantification of markers associated with activation of cGAS-STING signaling pathway

We measured the concentrations of 1) c-GAS (Cell Signaling, Danvers, MA, USA), 2) p-STING (Cell Signaling), 3) phosphorylated TANK-binding kinase-1 (p-TBK-1), 4) phosphorylated interferon regulatory factor 3 (p-IRF3; ABclonal, Woburn, MA, USA), and 5) IFN-α (LS-BIO, Lynnwood, WA, USA) in hippocampal lysates of male and female mice [17]. Since the activation of the cGAS-STING pathway stimulates the JAK-STAT signaling [42], we also measured concentrations of the phosphorylated JAK1 and JAK2 and the phosphorylated STAT1 and STAT3 in the hippocampal lysates of male and female aged mice. In addition, we performed qRT-PCR to measure the expression of ISG genes *Irf1*, *Icam1, Cd47, Irf5, Alcam, Irf7, Ifnγ,* and *Tnf* in RNA samples isolated from hippocampus tissue of aged Veh and aged-EVs groups of male and female mice.

### 2.12. Depletion of miR-30e-3p and miR-181a-5p in hiPSC-NSCs via transfection with Antagomirs

Previous studies have suggested that miR-30e-3p can inhibit NLRP3, by binding to the 3’ untranslated region (3’UTR) of NLRP3 mRNA [43], and miR-181a-5p can inhibit STING by directly binding to its mRNA [44], resulting in decreased expression of NLRP3 and STING proteins, respectively. Since both miR-30e-3p and miR-181a-5p are enriched in hiPSC-NSC-EVs, we investigated whether these miRs play significant roles in hiPSC-NSC-EVs-mediated inhibition of the activation of NLRP3 inflammasome and c-GAS-STING signaling. We first generated hiPSC-NSC-EVs with diminished expression of miR-30e-3p or miR-181a-5p. For this, P11 hiPSC-NSCs with 60% confluency in 6-well plates were transfected with 100 nM miR-30e-3p or 181a-5p inhibitor (AUM Biotech, Philadelphia, PA, USA) using Lipofectamine™ 2,000 transfection reagent (Invitrogen) in Opti-MEM media (ThermoFisher Scientific). Four hours later, Opti-MEM media were replaced with the NSC media, and cultures were maintained for an additional 48 hours with fresh NSC media. Next, spent media from cultures of transfected hiPSC-NSCs and their non-transfected counterparts were used to isolate EVs, as described in our earlier report [28], and qRT-PCR was employed to measure the levels of miR-30e-3p or miR-181a-5p. For this, total RNA from NSC-EVs was first isolated using the SeraMir Exosome RNA amplification kit (System Biosciences), and the miRCURY LNA RT Kit (Qiagen) was then used to convert 5 ng/μl of total RNA into cDNA. Next, miRCURY LNA miRNA SYBR Green PCR kit (Qiagen) and miRCURY LNA miRNA PCR assay primer mix (Qiagen) were employed to measure miRNA levels. Later, significantly depleted miR-30e-3p or miR-181a-5p levels in EVs generated from transfected hiPSC-NSCs were confirmed by comparing them with levels in EVs from non-transfected hiPSC-NSCs.

### 2.13. Assessing the effects of hiPSC-NSC-EVs with depleted miR-30e-3p on NLRP3 Inflammasome activation

To investigate the mechanism by which hiPSC-NSC-EVs suppress NLRP3 inflammasome activation, we performed an in vitro assay using RAW-ASC cells (InvivoGen, San Diego, CA, USA). These cells, derived from the murine RAW 264.7 macrophage line that are naturally ASC-deficient, were stably transfected with the murine ASC gene. RAW-ASC cells secrete IL-1β upon activation of canonical (e.g., NLRP3) or non-canonical (e.g., caspase-11) inflammasomes. For this, we first induced NLRP3 activation using nigericin, a well-characterized ionophore that promotes NLRP3 assembly with ASC adaptor protein and pro-caspase-1 to form an inflammasome complex. We next measured the concentrations of end products of NLRP3 inflammasome activation, the proinflammatory cytokines IL-1β and IL-18 in cells and culture media. Briefly, RAW-ASC cells were seeded at a density of 2 x 10^5^ cells/well in a 24-well plate and cultured for 24 hours in DMEM supplemented with 10% FBS and phenol red. Next, the media was replaced with serum-free DMEM containing 10 µM nigericin. Four hours later, the RAW ASC cultures were treated with 0.6 x 10^9^ hiPSC-NSC-EVs depleted with miR-30e-3p or naive hiPSC-NSC-EVs (n=3 wells/group). After 20 hours of treatment, the culture media and cells were collected to measure IL-1β and IL-18 levels using ELISA kits (R&D Systems).

### 2.14. Assessing the effects of hiPSC-NSC-EVs with depleted miR-181a-5p on STING activation

To understand the mechanism through which hiPSC-NSC-EVs reduce STING activation, we performed an in vitro assay using RAW-Lucia™ ISG cells (InvivoGen, San Diego, CA, USA). RAW-Lucia™ ISG cells express multiple pattern recognition receptors (PRRs), including the cyclic dinucleotide sensor STING. Activation of PRRs induces production of type 1 IFNs through the TBK-1-IRF3 axis. RAW-Lucia™ ISG cells also express the secreted Lucia luciferase reporter gene under the control of an ISG54 minimal promoter in conjunction with five IFN-stimulated response elements (ISRE). Hence, IRF3 activation in these cells can be determined by measuring the activity of the Lucia luciferase reporter. In this experiment, we used 2’,3’ cGAMP, which polymerized STING, and led to the activation of the TBK-1-IRF3 axis and the production of type 1 IFNs. We assessed IRF3 activation by measuring luciferase activity in the Lucia reporter via luminescence and IFNα production by ELISA. Briefly, RAW-LuciaTM-ISG cells were seeded in a 24-well plate at a density of 2 × 10^5^ cells/well in DMEM medium containing 10% FBS. After 24 hours, the media was replaced with serum-free DMEM (phenol red-free) containing 5 μM 2ʹ,3ʹ-cGAMP to activate STING. Four hours later, cultures were incubated with 0.6 x 10^9^ hiPSC-NSC-EVs depleted with miR-181a-5p or naive hiPSC-NSC-EVs (n=6 wells/group). Twenty hours later, supernatants from cultures were collected for measurement of luciferase activity, and cells were processed for IFNα quantification by ELISA (Ray Biotech). Lucia luciferase activity was measured from each culture by mixing 20µl of the culture supernatant with 50µl of the QANTI-LucTM 4 (1X) reagent in a 96-well white (opaque) plate. After gentle tapping, the luminescence was immediately recorded using a luminometer. The relative luminescence values were compared across the experimental groups.

### 2.15. Single-cell RNA sequencing (scRNA-seq) of microglia

Late middle-aged male mice (18 months old) were euthanized 7 days after administration of 12 × 10^9^ hiPSC-NSC-EVs or Veh (n=1/group). Fresh brains were immediately harvested and processed for the isolation of live microglia. Brain tissues were dissociated into single-cell suspensions using the gentleMACS Tissue Dissociator (Miltenyi Biotec, Gaithersburg, MD, USA). Microglia were isolated via magnetic-activated cell sorting (MACS) using CD11b-conjugated microbeads (Miltenyi Biotec) and MACS

Separators. Following live cell analysis and quality control measures, the live microglia were processed for scRNA-seq using the 10x Genomics Chromium GEM-X platform, targeting approximately 10,000 microglia per sample. Library preparation and sequencing were performed at the Texas A&M Institute for Genome Sciences and Society (TIGSS). Individually barcoded libraries were pooled and sequenced on a NextSeq 2000 system using a P4 100-cycle flow cell (Illumina, San Diego, CA, USA) according to the manufacturer’s instructions. The differentially expressed gene analysis was performed using scGEAToolbox [45]. The detailed methods for sequence alignment, filtering, and enrichment pathway analysis are provided in the supplementary file.

Next, we used the SelectCellByClass function in scGEAToolbox to extract microglial clusters from gene expression profiles in the SingleCellExperiment (SCE) object. The resulting cluster-specific SCE objects were then subjected to differential gene expression analysis, with detailed analysis provided in the supplementary materials. Furthermore, we performed GeneWalk analysis using the differentially expressed genes (DEGs) between the aged-Veh and aged-EVs groups. GeneWalk identifies relevant biological functions for genes by performing random walks on a knowledge graph that integrates gene networks with Gene Ontology (GO) annotations [46]. Mouse genes were first mapped to human orthologs, and all reactions between these orthologs were extracted from the knowledge base and automatically assembled into a gene network. Then, GO and annotations were added to these networks, resulting in the complete GeneWalk network. GeneWalk automatically identifies regulator genes (those with high connectivity to other input genes and a high fraction of relevant GO annotations) and moonlighting genes (those with many GO annotations but a low fraction of relevant ones). Moreover, since GeneWalk uses an adjusted p-value <0.1 for these classifications, we further filtered the results to retain only genes with an adjusted p-value <0.055. To further refine the set of strong regulator candidates, we applied two additional criteria: a fraction of relevant GO terms ≥0.7 and gene connectivity ≥75 (a connectivity of 100+ indicates a hub gene). Application of these filters excluded all moonlighting genes and a few regulator genes. We also removed GO terms in the cellular component domain. Finally, a GeneWalk scatterplot was generated that retained all genes with <250 connections.

### 2.16. Power analyses and statistics

The numbers of animals per group for neurobehavioral and brain tissue studies were determined via power analysis using data from our previous studies, the details of which are provided in the supplemental file. Statistical analyses were done using Prism software version 10.2. Within-group comparisons in neurobehavioral tests and two-group comparisons utilized either a two-tailed, unpaired Student’s t-test or a Mann-Whitney U-test, depending on whether the datasets had significantly different standard deviations. The variability across individual animals was comparable between the Aged-Veh and Aged-EVs groups, as indicated by the normality test comparing standard deviations between groups. A p-value of less than 0.05 was considered statistically significant in all tests. A two-way ANOVA followed by Tukey’s multiple comparisons post-hoc tests was employed to assess sex-dependent effects and the interaction between sex and hiPSC-NSC-EVs treatment.

## 3. RESULTS

### 3.1. Attributes of hiPSC-NSC-EVs employed in the study

The P11 hiPSC-NSC cultures employed in the study exclusively comprised NSCs, as all cells expressed the NSC markers nestin and Sox-2 (Fig. 1 [A]). EVs isolated from these cultures using anion-exchange and size-exclusion chromatography had a mean size of 130.9 nm (Fig. 1 [B]). These EVs exhibited specific markers, such as the tetraspanins CD63 and CD81, and ALIX. However, they did not contain the cytoplasmic marker calnexin, which is present in hiPSC-NSCs (Fig. 1 [C] and Suppl. Fig. 1). Additionally, transmission electron microscopy confirmed the presence of double-membrane-bound vesicles in the EV preparations (Fig. 1 [D]). Our previous study demonstrated that various protein and miRNA cargoes carried by these hiPSC-NSC-derived EVs exhibit antioxidant, anti-inflammatory, and neuroprotective properties [26, 28].

### 3.2. Biodistribution of IN-administered hiPSC-NSC-EVs in the brain of late middle-aged mice

The biodistribution of IN-administered PKH-26-labeled hiPSC-NSC-EVs was examined in multiple brain regions using markers of neural cells at 6 hours post-administration (Fig. 1 [E-F]) in both male and female mice. Such analysis revealed that hiPSC-NSC-EVs permeated virtually all brain regions within six hours post-administration, as demonstrated in our earlier studies for naïve mice [26] and 5x Familial AD (5xFAD) mice [27]. In all brain regions, EVs were taken up by microglia and neurons. EVs were also seen interacting with the cell membranes of astrocytes and oligodendrocytes. As this study focused on hiPSC-NSC-EVs treatment-mediated changes in microglia and astrocytes, examples of hiPSC-NSC-EVs incorporation by IBA-1+ microglia and the interaction of hiPSC-NSC-EVs with the plasma membrane of soma and processes of GFAP+ astrocytes in the hippocampal subregions (DG, CA1, and CA3) are illustrated for both male and female mice (Fig. 1 [E-F]). hiPSC-NSC-EVs incorporation by IBA-1+ microglia and the interaction of hiPSC-NSC-EVs with GFAP+ astrocytes in other brain regions are illustrated in the Supplemental File (Suppl. Fig. 2). Overall, the biodistribution of IN-administered hiPSC-NSC-EVs in late middle-aged mice mirrored the distribution pattern observed in young adult mice [26–27].

### 3.3. hiPSC-NSC-EVs reduced astrocyte hypertrophy in the hippocampus

An evaluation of the morphology of GFAP+ astrocytes in the hippocampus suggested that hypertrophy of astrocytes decreased in the aged-EVs group compared to the aged-Veh group.

Representative images from the CA3 subfield of the hippocampus from male and female aged-Veh and aged-EVs groups are illustrated (Fig. 2 [A-D]). Measurements of the area fraction of GFAP+ astrocytic elements in the hippocampus, using ImageJ, confirmed that both male and female subjects in the aged-EVs group exhibited lower levels of GFAP+ structures than their counterparts in the aged-Veh group (p<0.0001; Fig. 2 [E-F]). A two-way ANOVA revealed sex-dependent differences in astrocyte hypertrophy within the aged-Veh group, with male mice exhibiting higher levels than female mice (p<0.05). However, an analysis of the interaction between sex and EVs treatment demonstrated that both male and female mice responded positively to EVs treatment (p<0.05; Suppl. Table 2). Thus, treatment with hiPSC-NSC-EVs in late middle age can reduce astrocyte hypertrophy in the aged hippocampus.

**Figure 2:**
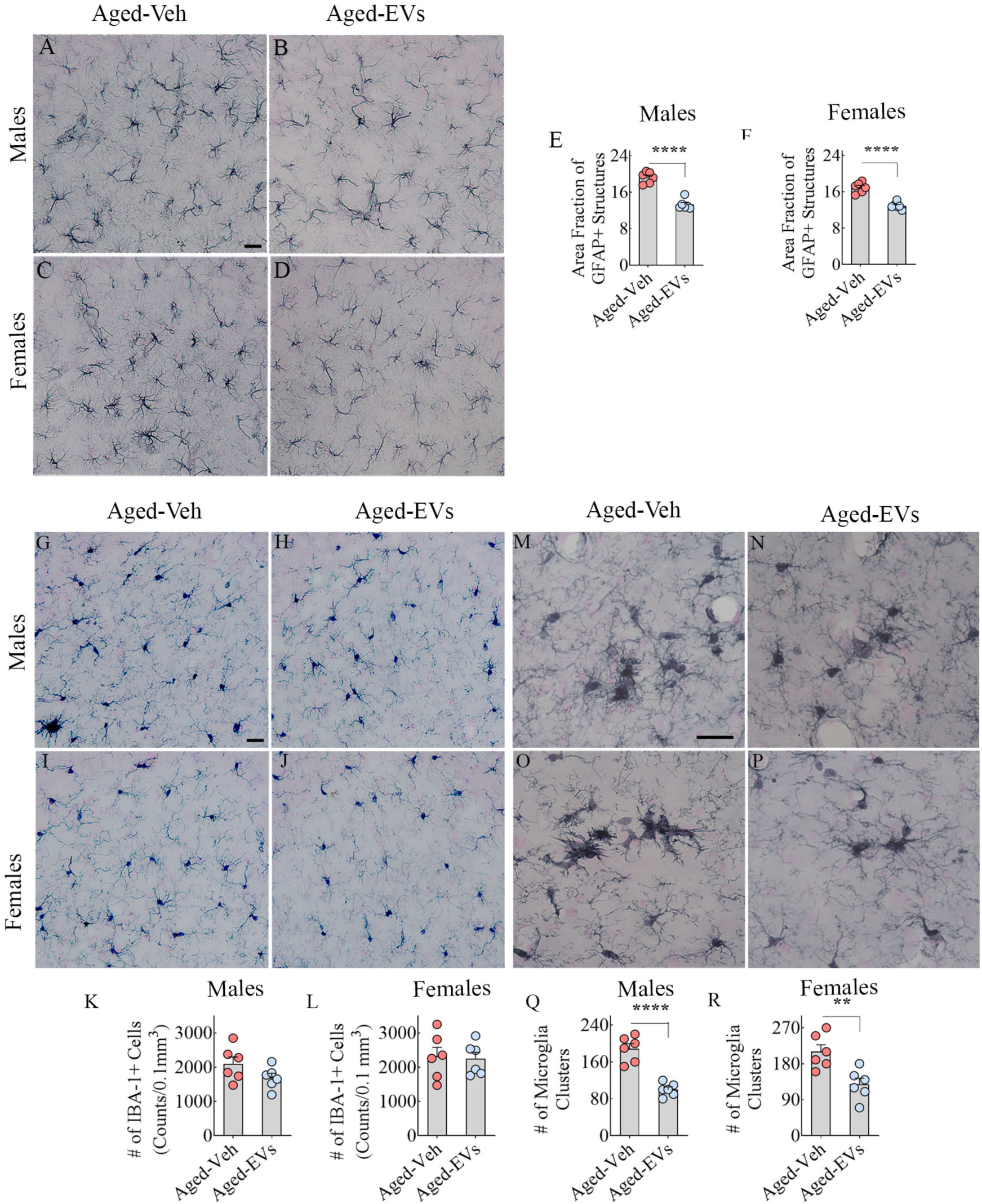
Intranasal administration of extracellular vesicles from human induced pluripotent stem cell-derived neural stem cells (hiPSC-NSC-EVs) to late middle-aged mice reduced hypertrophy of astrocytes and microglial clusters. Figures A-D illustrate the representative images of GFAP+ astrocytes from aged-Veh (A, C) and aged-EVs (B, D) groups for males (A-B) and females (C-D). Bar charts E and F compare the area fraction (AF) of GFAP+ structures in the hippocampus of males (E) and females (F) between the aged-Veh and aged-EV groups. Figures G-J illustrate the representative images of IBA-1+ microglia from aged-Veh (G, I) and aged-EVs (H, J) groups for males (G-H) and females (I-J). Bar charts K and L compare the density of IBA-1+ microglia per 0.1 mm^3^ in the hippocampus in males (K) and females (L) between the aged-Veh and aged-EV groups. Figures M-P illustrate the representative images of microglia clusters from aged-Veh (M, O) and aged-EVs (N, P) groups for males (M-N) and females (O-P). Bar charts Q-R compare the number of microglia clusters in the hippocampus of males (Q) and females (R) between the aged-Veh and aged-EV groups. Scale bar, A-D, G-J, M-P = 25 μm **, p<0.01; and ****, p<0.0001.

### 3.4. hiPSC-NSC-EVs reduced microglial numbers and clusters in the hippocampus

Analysis of the distribution of IBA-1+ microglia in the hippocampus suggested a comparable density of microglia in the aged-EVs group compared to the aged-Veh group. Representative images from the CA3 subfield of the hippocampus from male and female aged-Veh and aged-EVs groups are illustrated (Fig 2 [G-J]). Stereological quantification of the microglial numbers per unit volume of the hippocampus confirmed that both male and female mice from the aged-Veh and aged-EVs group displayed comparable densities of IBA-1+ microglia in the hippocampus (p>0.05; Fig. 2 [K-L]), implying that EVs treatment did not alter the microglial number. However, an examination of microglial clusters comprising hypertrophied microglia, indicative of disease-associated microglial phenotypes [47–48], in the hippocampus revealed lower levels in the aged-EVs group than in the aged-Veh group.

Representative examples from the dentate molecular layer of male and female mice in the aged-Veh and aged-EVs groups are shown (Fig. 2 [M-P]). Quantification of these microglial clusters along the septo-temporal axis of the hippocampus confirmed that both male and female mice in the aged-EVs group had fewer microglial clusters compared to those in the aged-Veh group (p<0.01-0.0001, Fig. 2 [Q-R]). A two-way ANOVA analysis revealed neither sex-dependent differences in microglial numbers or clusters in aged-Veh and aged-EVs groups nor interactions between sex and hiPSC-NSC-EVs treatment (Suppl. Table 2). Thus, hiPSC-NSC-EVs treatment in late middle age can reduce disease-associated microglia phenotypes in the aged hippocampus.

### 3.5. hiPSC-NSC-EVs reduced oxidative stress with enhanced NRF2 in the hippocampus

Proficiency of hiPSC-NSC-EVs to reduce oxidative stress was investigated by measuring markers of oxidative stress (MDA and PCs), the master regulator of oxidative stress (NRF2), and antioxidants (SOD and CAT). Compared to the aged-Veh group, both males and females in the aged-EVs group displayed reduced concentrations of MDA and PCs (p<0.01-0.0001, Fig. 3 [A-B, F-G]), increased concentrations of NRF-2 and SOD (p<0.01, Fig. 3 [C-D, H-I]). The concentration of CAT in the aged-EVs group was increased in females (p<0.01, Fig. 3 [J]) but not in males (p>0.05, Fig. 3 [E]). Two-way ANOVA analyses showed neither sex-dependent differences in the concentrations of MDA, PCs, NRF2, and SOD in aged-Veh and aged-EVs groups nor interactions between sex and hiPSC-NSC-EVs treatment. However, CAT concentration was sex-dependent in the aged-EVs group, with females exhibiting higher CAT concentration (p<0.01; Suppl Table 2), and the interaction between sex and EVs treatment revealed that only female mice responded positively to EVs treatment (p<0.01; Suppl Table 2). Thus, hiPSC-NSC-EVs treatment in late middle age can reduce oxidative stress in the aged hippocampus.

**Figure 3:**
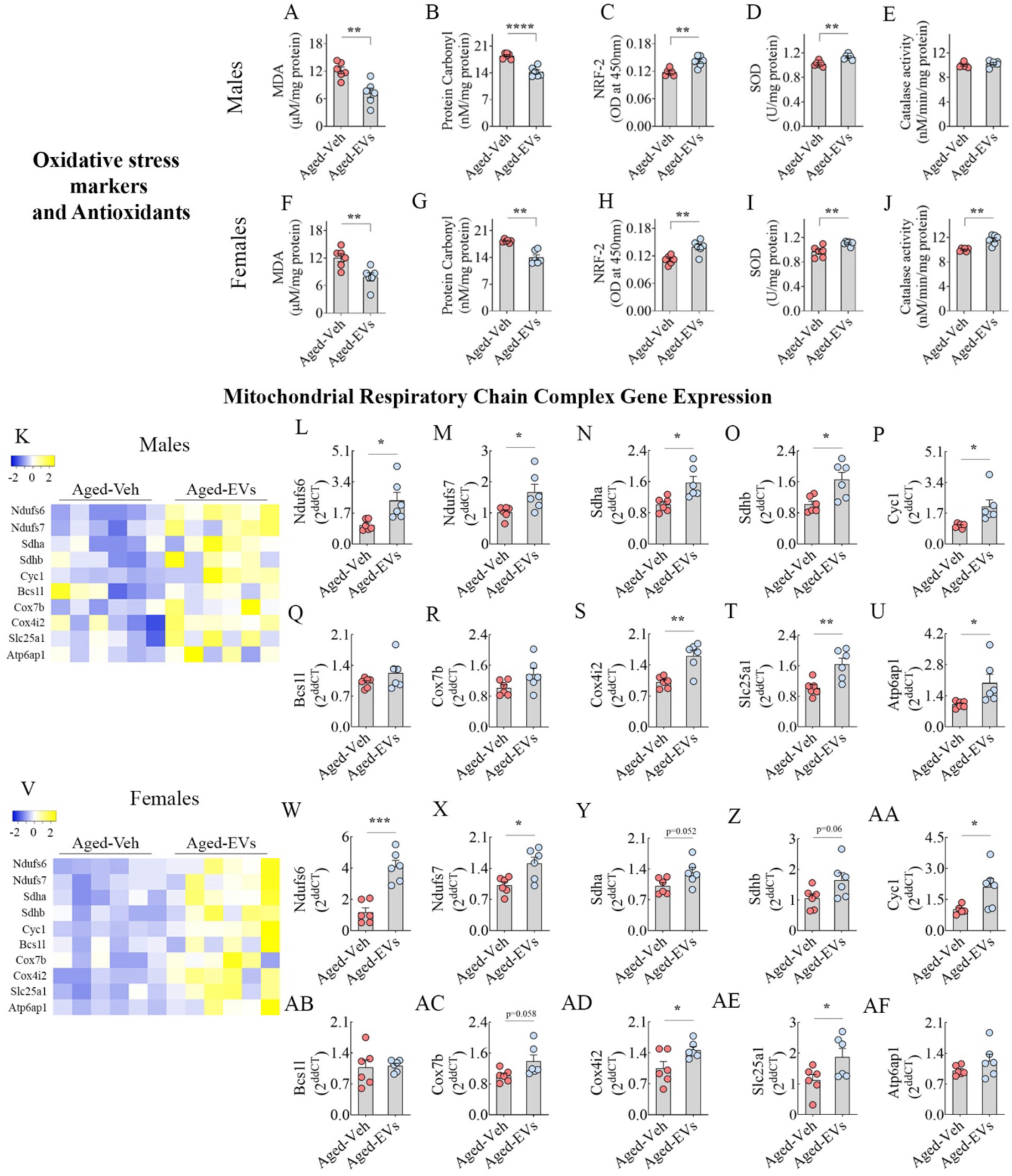
Intranasal administration of extracellular vesicles from human induced pluripotent stem cell-derived neural stem cells (hiPSC-NSC-EVs) to late middle-aged mice reduced oxidative stress and ***normalized the mitochondrial respiratory chain gene expression.*** The bar charts A-J compare malondialdehyde (A, F), protein carbonyl (B, G), NRF-2 (C, H), SOD (D, I), and catalase (E, J) between aged-Veh and aged-EVs groups in males (A-E) and females (F-J). Heatmaps K and V compare the expression of multiple mitochondrial respiratory chain genes between aged-Veh and aged-EV groups in male (K) and female (V) mice. Bar charts L-U (male), and W-AF (female) show the relative expression of genes Ndufs6 (L, W), Ndufs7 (M, X), Sdha (N, Y), Sdhb (O, Z), Cyc1 (P, AA), Bcs1l (Q, AB), Cox7b (R, AC), Cox4i2 (S, AD), Slc25a1 (T, AE) and Atp6ap1 (U, AF) in aged-Veh and aged-EV groups. *, p<0.05; **, p<0.01; ***, p<0.001; ****, p<0.0001.

### 3.6. hiPSC-NSC-EVs enhanced mitochondrial respiratory chain gene expression in the hippocampus

The competence of hiPSC-NSC-EVs to enhance the expression of genes linked to the mitochondrial respiratory chain was investigated in the hippocampus, as increased oxidative stress damages mitochondria, thereby impairing mitochondrial respiratory chain activity [49]. The measured genes comprised those linked to the activity of complex I *(Ndufs6, Ndufs7)*, complex II *(Sdha, Sdhb)*, complex III *(Cyc1, Bcs1l)*, complex IV *(Cox7b, Cox4i2)*, and complex V *(Slc25a1, Atp6ap1)* (Fig. 3 [K, V]). Compared to the aged-Veh group, males in the aged-EVs group displayed significantly increased expression of many genes, including *Ndufs6, Ndufs7, Sdha, Sdhb, Cyc1, Cox4i2, Slc25a1,* and *Atp6ap1* (p<0.05-0.01, Fig. 3 [L-P, S-U]). Females in the aged-EVs groups showed a similar trend but with fewer genes *(Ndufs6, Ndufs7, Cyc1, Cox4i2, Slc25a1)* showing significantly increased expression than their counterparts in the aged-Veh group (p<0.05-0.001, Fig. 3 [W-X, AA, AD-AE]). Two-way ANOVA analyses showed neither sex-dependent differences in aged-Veh and aged-EVs groups nor interactions between sex and hiPSC-NSC-EVs treatment for most of the measured genes linked to the mitochondrial respiratory chain (p>0.05, Suppl. Table 2). However, the expression of *Ndufs6* was sex-dependent in the aged-EVs group, with females exhibiting an increased expression (p<0.05, Suppl Table 1), but analysis of the interaction between sex and EVs treatment revealed that *Ndufs6* expression in both male and female mice responded positively to EVs treatment (p<0.01; Suppl Table 2). Thus, hiPSC-NSC-EVs treatment in late middle-aged male and female mice enhanced mitochondrial respiratory chain gene expression in the aged hippocampus.

### 3.7. hiPSC-NSC-EVs restrained the activation of NLRP3 inflammasome cascade in the hippocampus

Proficiency of hiPSC-NSC-EVs to restrain the activation of NLRP3 inflammasomes was examined by three measures, which include quantification of 1) the expression of associated genes such as *Nlrp3, Pycard, Casp1, Il1β, Il18* (Fig. 4 [A-L]), 2) percentages of microglia exhibiting NLRP3-ASC complexes (Fig. 4 [M-Z]), and 3) the concentrations of proteins involved in their activation, including NF-kB-p65, NLRP3, ASC, Cleaved Caspase-1, IL-18, and IL-1β, (Fig. 4 [AA-AL]). Compared with the aged-Veh group, males in the aged-EVs group showed decreased expression of all measured genes, with *Nlrp3, Pycard,* and *Il18* showing significant declines (p<0.05-0.001; Fig. 4 [B-C, E]). Females in the aged-EVs groups showed a similar trend, with *Nlrp3* and *il18* showing significantly decreased expression compared to their counterparts in the aged-Veh group (p<0.05, Fig. 4 [H, K]). Next, the extent of NLRP3-ASC inflammasome complexes within microglia of the hippocampus was examined via triple immunofluorescence for NLRP3, ASC, and IBA-1 and Z-section analysis (Fig. 4 [M-Y]). Compared with the aged-Veh group, both males and females in the aged-EVs group showed lower percentages of microglia expressing NLRP3-ASC inflammasome complexes (p<0.01, Fig. 4 [S, Z]).

**Figure 4:**
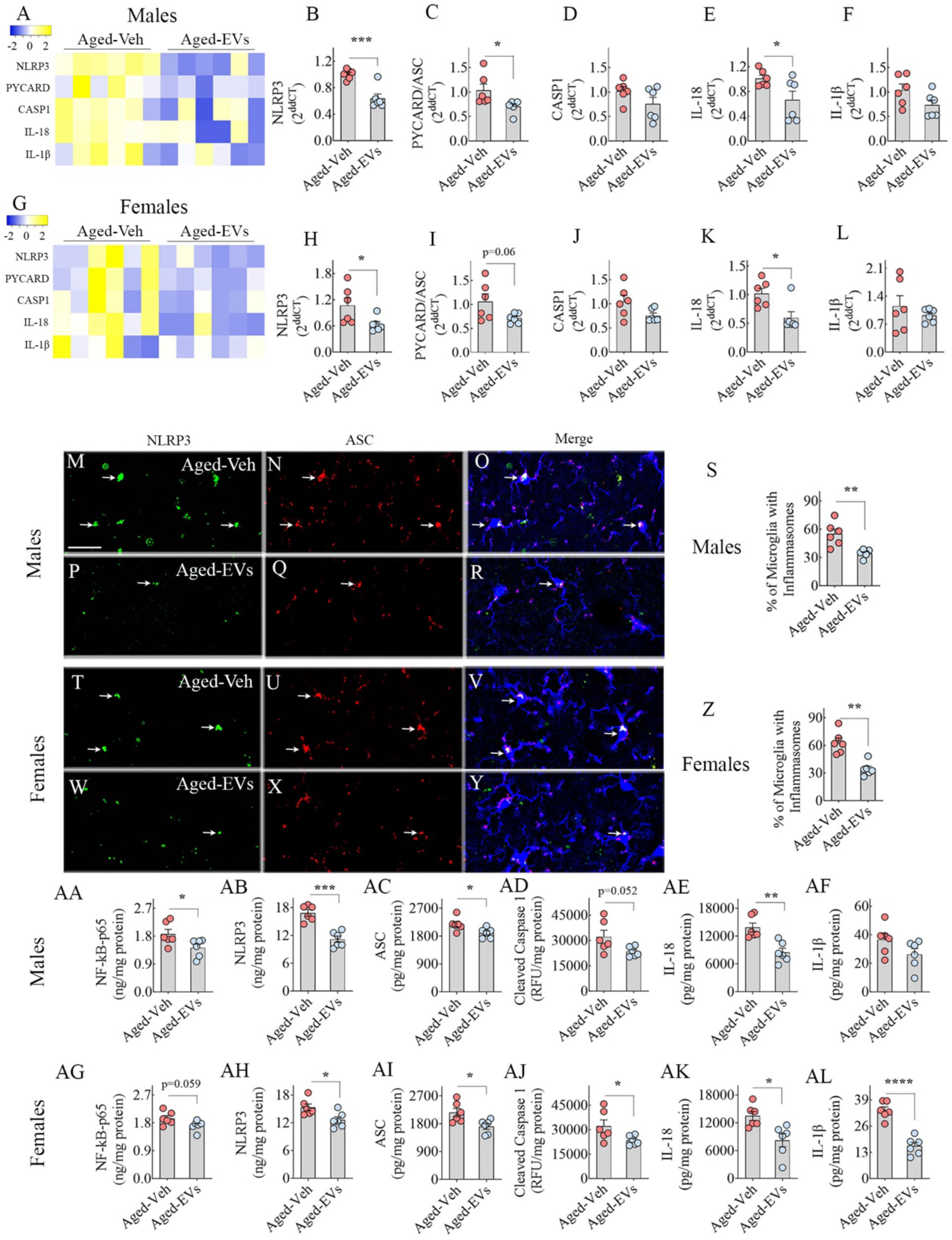
Intranasal administration of extracellular vesicles from human induced pluripotent stem cell-derived neural stem cells (hiPSC-NSC-EVs) to late middle-aged mice inhibited NOD-, LRR-, and pyrin domain-containing protein 3 (NLRP3) and apoptosis-associated speck-like protein containing a CARD (ASC) inflammasome complex formation. Heatmaps A and G compare the expression of multiple NLRP3 inflammasome genes between aged-Veh and aged-EV groups in male (A) and female (G) mice. The bar charts B-F, H-L compare NLRP3 (B, H), PYCARD (C, I), CASP1 (D, J), IL-18 (E, K), and IL-1β (F, L) across aged-Veh and aged-EVs groups in males (B-F) and females (H-L). Figures M-R and T-Y illustrate examples of NLRP3 inflammasome complexes co-expressing NLRP3 (green) and ASC (red) in IBA-1+ microglia (blue) in the CA3 subfield of the hippocampus from male (M-R) and female (T-Y) mice belonging to aged-Veh (M-O, T-V) and aged-EVs (P-R, W-Y) groups. The bar charts S and Z compare the percentages of microglia with inflammasome complexes in males (S) and females (Z). The bar graphs AA-AL compare the concentrations of different proteins involved in the activation of NLRP3 inflammasomes, such as NF-kB p65 (AA, AG), NLRP3 (AB, AH), ASC (AC, AI), cleaved caspase-1 (AD, AJ), IL-18 (AE, AK), and IL-1β (AF, AL) between aged-Veh and aged-EVs groups of male (AA-AF) and female (AG-AL) mice. M-Y = 25 μm; *, p<0.05; **, p<0.01; ***, p<0.001; ****, p<0.0001.

To corroborate the above findings, the concentrations of proteins involved in NLRP3 inflammasome activation were measured. Compared with the aged-Veh group, males in the aged-EVs group showed decreased concentrations of all measured proteins, with NF-kB-p65, NLRP3, ASC, and IL-18 showing significant declines (p<0.05-0.001; Fig. 4 [AA-AC, AE]). Females in the aged-EVs groups showed a similar trend with NLRP3, ASC, Cleaved Caspase-1, IL-18, and IL-1β showing significantly decreased concentrations than their counterparts in the aged-Veh group (p<0.05-0.0001; Fig. 4 [AH-AL]). Two-way ANOVA analyses showed neither sex-dependent differences in Aged-Veh and aged-EVs groups nor interactions between sex and hiPSC-NSC-EVs treatment for all measured genes and a vast majority of the measured proteins linked to NLRP3 inflammasome activation (p>0.05; Suppl. Table 2). The exception is the concentration of NLRP3, which showed an interaction between sex and EVs treatment, although both males and females responded positively to EVs treatment (p<0.05; Suppl Table 2). Thus, hiPSC-NSC-EV treatment in late middle-aged male and female mice significantly inhibited NLRP3 inflammasome activation in the aged hippocampus. Such effect was evident from reduced concentrations of 1) the transcription factor NF-kB-p65 that triggers the formation of NLRP3 inflammasome complexes, 2) proteins that facilitate their activation (NLRP3, ASC, and Cleaved Caspase-1), and 3) end products of NLRP3 inflammasome activation (IL-1β and IL-18).

### 3.8. hiPSC-NSC-EVs diminished the activation of p38/MAPK signaling in the hippocampus

Since the end products of NLRP3 inflammasome activation (IL-1β and IL-18) and increased oxidative stress can mediate the downstream activation of p38/MAPK inflammatory signaling cascade (Fig. 5 [A]), the proficiency of hiPSC-NSC-EVs to reduce the concentrations of proteins involved in p38/MAPK signaling activation (Myd88, Ras, pMAPK, and AP-1) and some of the end products of p38/MAPK activation (TNF-α and IL-18) were measured (Fig. 5 [B-M]). In comparison to the aged-Veh group, both males and females in the aged-EVs group displayed decreased concentrations of all measured proteins, with Myd88, Ras, pMAPK, AP-1, and TNF-α showing a significant decline in both sexes (p<0.05-0.001; Fig. 5 [B-F, H-L]). The concentration of IL-8 was significantly decreased in females (p<0.05; Fig. 5 [M]) but showed only a trend in males (p=0.05; Fig. 5 [G]). Two-way ANOVA analyses did not show sex-dependent differences in aged-Veh and aged-EVs groups for the concentrations of Myd88, Ras, AP-1, and IL-8 (p>0.05; Suppl. Table 2). The concentration of pMAPK showed sex-specific differences, with higher levels in females in the aged-EVs group (p<0.05; Suppl Table 2). The concentration of TNF-α also showed sex-specific differences, with higher levels in the hippocampus of males within the aged-EVs group. However, post-hoc tests did not reveal significant differences (Suppl Table 2). Furthermore, no interaction was found between sex and hiPSC-NSC-EVs treatment for any of the proteins linked to p38/MAPK activation (p>0.05; Suppl Table 2). Thus, hiPSC-NSC-EV treatment in late middle-aged male and female mice prevented p38/MAPK inflammatory signaling activation in the aged hippocampus.

**Figure 5:**
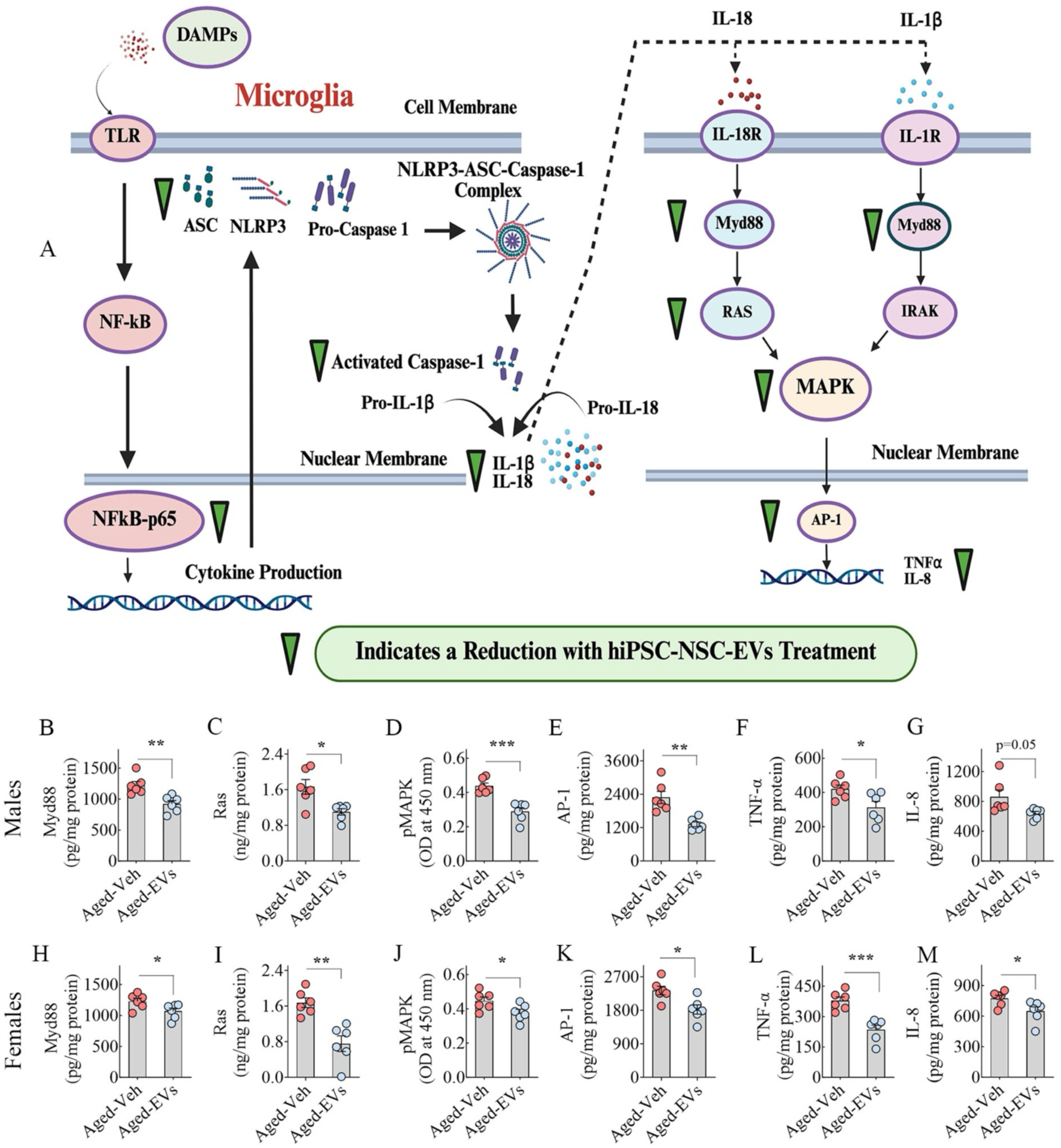
Intranasal administration of extracellular vesicles from human induced pluripotent stem cell-derived neural stem cells (hiPSC-NSC-EVs) to late middle-aged mice inhibited p38/mitogen-activated protein kinase (p38/MAPK) signaling. Figure A is a cartoon depicting the various steps in NLRP3 inflammasome activation and how its end products (IL-18 and IL-1β) activate p38/MAPK signaling, leading to the increased production of proinflammatory cytokines in the aged brain. The downward-pointing green arrowheads depict various proteins that are reduced in concentration with hiPSC-NSC-EVs treatment. The bar charts in B-M compare the concentrations of p38MAPK signaling-related proteins between aged-Veh and aged-EVs groups in male and female mice. The proteins include Myd88 (B, H), Ras (C, I), pMAPK (D, J), transcription factor AP-1 (E, K), and end products such as TNF-α (F, L) and IL-8 (G, M) in male (B-G) and female (H-M) mice. *, p<0.05; **, p<0.01; ***, p<0.001.

### 3.9. hiPSC-NSC-EVs curtailed the activation of cGAS-STING and the downstream JAK-STAT signaling and ISG expression in the hippocampus

As the cGAS–STING pathway and downstream JAK-STAT pathway (Figure 6 [A]) is one of the drivers of aging-related inflammation in both peripheral organs and the brain [14], the ability of hiPSC-NSC-EVs to reduce the concentrations of proteins involved in cGAS-STING signaling activation (p-cGAS, p-STING, p-TBK1 and p-IRF3) and one of its end products (IFN-α) was measured (Fig. 6 [B-K]). Compared with the aged-Veh group, both males and females in the aged-EVs group showed decreased concentrations of all measured proteins, with p-cGAS, p-STING, and p-IRF3 showing significant declines in both sexes (p<0.05-0.01; Fig. 6 [B-C, E, G-H, J]). The concentration of p-TBK1 was significantly decreased in females (p<0.0001, Fig. 6 [I]) but showed only a trend in males (p=0.058; Fig. 6 [D]). The concentration of IFN-α was significantly decreased in males (p<0.001, Fig. 6 [F]) but showed only a trend in females (p=0.0509, Fig. 6 [K]). Two-way ANOVA analyses did not show sex-dependent differences in aged-Veh and aged-EVs groups for the concentrations of p-cGAS, p-STING, p-TBK1, and IFN-α (p>0.05, Suppl. Table 2). The concentration of p-IRF3 showed sex-specific differences, with higher levels in females in the aged-EVs group (p<0.05, Suppl Table 2). Furthermore, no interaction was found between sex and hiPSC-NSC-EVs treatment for most of the proteins linked to cGAS-STING activation (p>0.05; Suppl Table 2). The exception is the concentration of IFN-α, which showed an interaction between sex and EVs treatment, although both males and females responded positively to EVs treatment (p<0.05; Suppl Table 2). Thus, hiPSC-NSC-EVs treatment in late middle age considerably diminished p38/MAPK inflammatory signaling activation in the hippocampus of both aged male and female mice.

**Figure 6:**
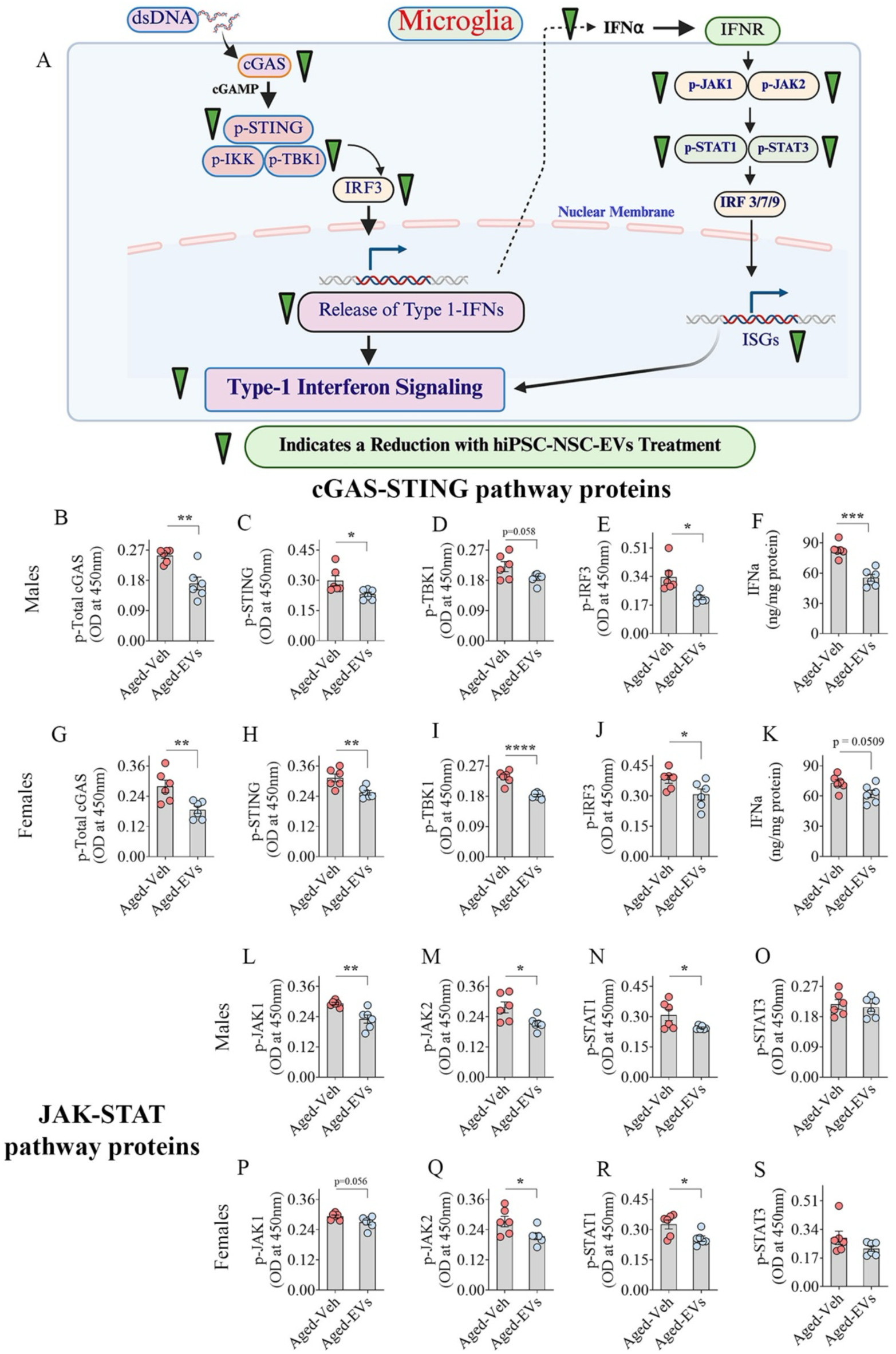
Intranasal administration of extracellular vesicles from human induced pluripotent stem cell-derived neural stem cells (hiPSC-NSC-EVs) to late middle-aged mice inhibited cGAS-STING and JAK-STAT pathway activation. Figure A is a cartoon depicting the various steps in the activation of cGAS-STING-IFN-1 signaling in the aged brain. The downward-pointing green arrowheads depict various proteins in this signaling pathway that are reduced in concentration following hiPSC-NSC-EV treatment. The bar charts B-K compare the concentration of proteins linked to the cGAS-STING pathway, such as p-total cGAS (B, G), p-STING (C, H), p-TBK1 (D, I), p-IRF3 (E, J), and IFN-α (F, K) between Aged-Veh and Aged-EVs groups in male (B-F) and female (G-K) mice. The bar charts L-S compare the concentrations of proteins linked to the JAK-STAT pathway, including p-JAK1 (L, P), p-JAK2 (M, Q), p-STAT1 (N, R), and p-STAT3 (O, S) between Aged-Veh and Aged-EVs groups in male (L-O) and female (P-S) mice. *, p<0.05; **, p<0.01; ***, p<0.001; ****, p<0.0001.

Since increased release of IFN-α can lead to the activation of the downstream IFN-1 signaling through the JAK-STAT protein activation [50], the proficiency of hiPSC-NSC-EVs to reduce the concentrations of proteins involved in JAK-STAT signaling (p-JAK1, p-JAK2, p-STAT1 and p-STAT3) was measured (Fig. 6 [L-S]). Compared to the aged-Veh group, both males and females in the aged-EVs group displayed significantly decreased concentrations of p-JAK2 and p-STAT1 (p<0.05, Fig. 6 [M-N, Q-R]). The concentration of p-JAK1 was significantly decreased in males (p<0.01, Fig. 6 [L]) but showed only a decreased trend in females (p=0.056, Fig. 6 [P]). The concentrations of p-STAT3 did not differ between aged-Veh and aged-EVs groups in both males and females (p>0.05, Fig. 6. [N, R]). Two-way ANOVA analyses showed neither sex-dependent differences in aged-Veh and aged-EVs groups nor interactions between sex and hiPSC-NSC-EVs treatment for all measured proteins linked to JAK-STAT signaling (p>0.05, Suppl. Table 2). Thus, hiPSC-NSC-EVs treatment in late middle age considerably diminished JAK-STAT activation involved in IFN-1 signaling in the hippocampus of both aged male and female mice.

Next, we confirmed decreased expression of select ISGs in the hippocampus of both males and females in the aged-EVs groups compared to their counterparts in the aged-Veh groups (Suppl. Fig. 3 [A-R]). In males receiving hiPSC-NSC-EVs, the expression of genes such as *Icam1, Cd47, Irf5, Irf7, IFNγ,* and *Tnf* was downregulated (p<0.05-0.01, Suppl. Fig. 3 [C-E, G-I]). Whereas in females receiving hiPSC-NSC-EVs, the expression of *Icam1, Cd47,* and *Irf7* was reduced (p<0.05-0.01, Suppl. Fig. 3 [L-M, P]).

### 3.10. hiPSC-NSC-EVs treatment in late middle-age led to improved cognitive function in old age

We first examined whether anxiety levels varied between animals belonging to the Aged-Veh and Aged-EVs groups using data from an OFT (Fig. 7 [A]). Comparison of time spent in the central region of the open field and the frequency of entries into the central region revealed similar behavior between aged-Veh and aged-EVs groups for both males and females (p>0.05, Figure 7 [B-E]). Thus, the results of cognitive and memory tests are not influenced by differences in anxiety levels between the aged-Veh and aged-EVs groups. We next investigated recognition and object-location memories using a NORT (Fig. 7 [F]) and an OLT (Fig. 7 [N]). In NORT, animals proficient in recognition memory prefer to explore the novel object (NO) over the familiar object (FO). In OLT, animals competent for object location memory prefer to explore the object in the novel place (OINP) over the object in the familiar place (OIFP). We initially confirmed that the male and female mice chosen for the study do display recognition and object-location memory impairments at the time point of hiPSC-NSC-EVs or Veh intervention. These results were obtained from separate cohorts of male and female mice to avoid the confounding effects of repeated behavioral testing on outcomes. These results are detailed in the Supplemental File (Suppl. Fig. 4).

**Figure 7:**
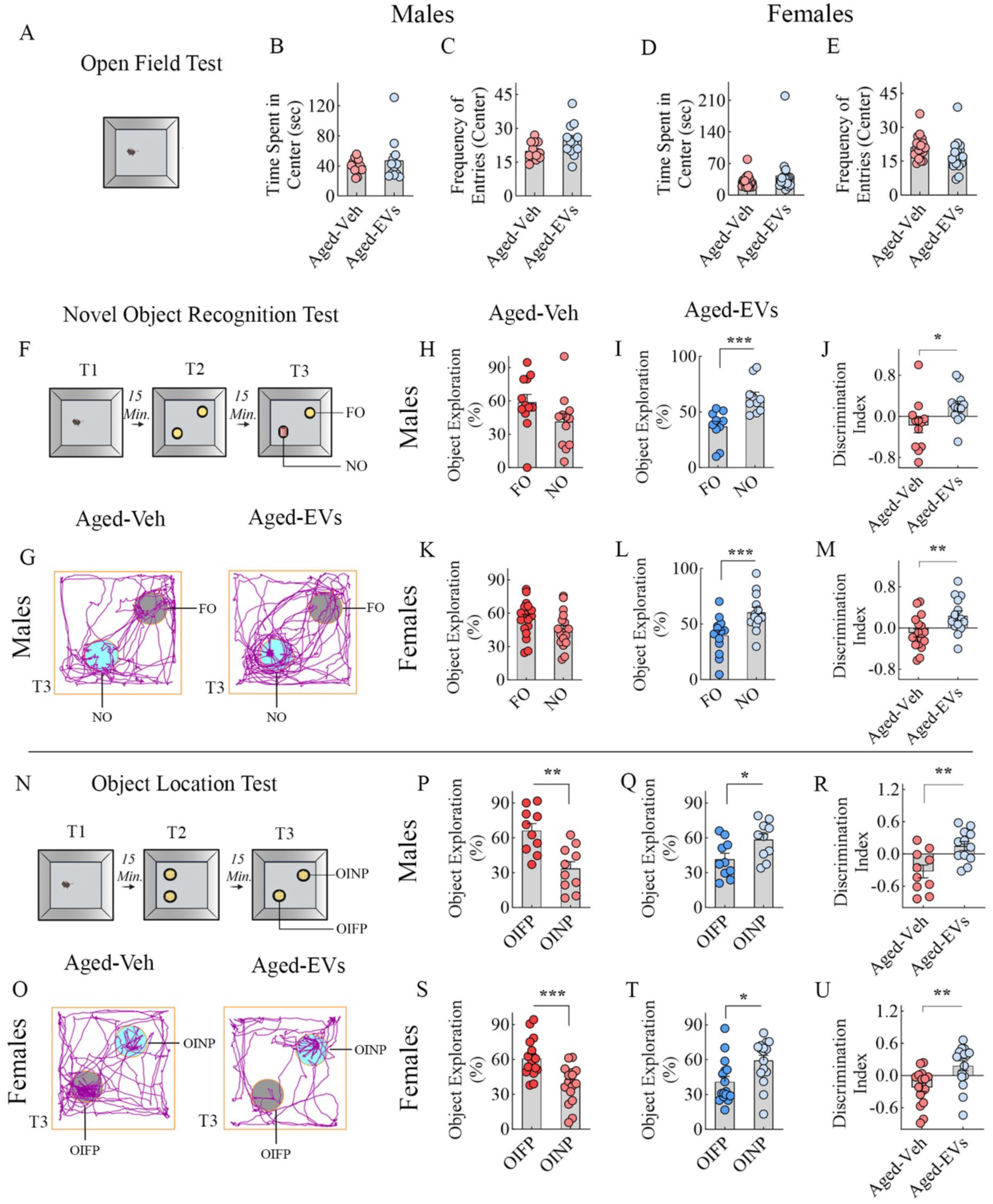
Intranasal administration of extracellular vesicles from human induced pluripotent stem cell-derived neural stem cells (hiPSC-NSC-EVs) to late middle-aged mice improved cognitive dysfunction. Cartoon A depicts the open field test (OFT). Bar chart B-E compares the time spent in the center (B, D) and frequency of entries to the center (C, E) in males (B-C) and females (D-E) between AD-Veh and AD-EVs groups. Cartoon F depicts different trials (T1-T3) in a novel object recognition test (NORT). Representative tracings in G illustrate the exploration of objects by male mice from the Aged-Veh and Aged-EVs groups in T3. The bar charts in H-I and K-L compare percentages of object exploration times spent with the familiar object (FO) vis-à-vis the novel object (NO) between Aged-Veh (H, K) and Aged-EV groups (I, L) for males (H-I) and females (K-L). The bar charts J and M compare the NO discrimination index values between Aged-Veh and Aged-EVs groups in males (J) and females (M). Cartoon N depicts different trials (T1-T3) in an object location test (OLT). Representative tracings in O illustrate the exploration of objects by female mice from aged-Veh and aged-EVs groups in T3. The bar charts in P-Q and S-T compare percentages of object exploration times spent with the object in the familiar place (OIFP) vis-à-vis the object in the novel place (OINP) between aged-Veh (P, S) and aged-EV groups (Q, T) in males (P-Q) and Females (S-T). The bar charts R and U compare the OINP discrimination index values between Aged-Veh and Aged-EVs groups in males (R) and females (U). *, p<0.05; **, p<0.01; ***, p<0.001.

We next examined cognitive and memory function in animals from the aged-Veh and aged-EVs groups between 19.5 and 20.5 months of age (i.e., commencing 1 month after the last Veh/EVs treatment). In NORT (Fig. 7 [F-M]), male and female mice in the aged-Veh group showed no preference to explore either the NO or the FO (p>0.05; Fig. 7 [H, K]), whereas male and female mice in the aged-EVs group preferred to explore the NO for longer durations than the FO (p<0.001; Fig. 7 [I, L]).

Representative tracings of mouse movement from the aged-Veh and aged-EVs groups while exploring the FO and NO during trial-3 (T3) of an NORT are shown (Fig. 7 [G]). In OLT (Fig. 7 [N-U]), mice in the aged-Veh group showed a predilection for exploring the OIFP over the OINP (p<0.01-0.001; Fig. 7 [P, S]), whereas mice in the aged-EVs group exhibited a preference to explore the OINP for longer durations than the OIFP (p<0.05; Fig. 7 [Q, T]). Representative tracings of mouse movement from the aged-Veh and aged-EVs groups while exploring the OINP and OIFP in T3 of an OLT are shown (Fig. 7 [O]). Thus, male and female mice in the aged-Veh group displayed impairments in recognition and object location memory formation. In contrast, male and female mice in the aged-EVs groups exhibited proficiency in recognition and object location memory. Such a conclusion is also confirmed by significantly lower NO/OINP-DI values in male and female mice in the aged-Veh group compared to their counterparts in the aged-EVs group (p<0.05-0.01; Fig. 7 [J, M, R, U]). The total object exploration times spent in trials 2 and 3 (T2 and T3) in NORT and OLT are presented in the supplemental file (Suppl. Fig. 5). Two-way ANOVA showed neither sex-dependent differences in aged-Veh and aged-EVs groups nor interactions between sex and hiPSC-NSC-EVs treatment for NO or OINP-DI values in both NORT and OLT (p>0.05; Suppl. Table 2).

### 3.11. Depletion of miR-30e-3p within hiPSC-NSC-EVs diminished their ability to inhibit NLRP3 Inflammasome activation

By employing RAW-ASC cells that are transfected with the murine ASC gene, we investigated how hiPSC-NSC-EVs inhibit nigericin-induced NLRP3 activation (Fig. 8 [A]). For this, we first confirmed significantly reduced miR-30e-3p levels in EVs isolated from hiPSC-NSCs transfected with antagomirs targeting miR-30e-3p-5p. EVs from transfected hiPSC-NSCs displayed a ∼79.61% decline in miR-30e-3p expression compared to EVs from naïve hiPSC-NSCs (Fig. 8 [B]). No such reduction was observed in EVs isolated from hiPSC-NSCs transfected with the scrambled miR-30e-3p-5p antagomir. A schematic representation of the experimental workflow is shown in Fig. 8 [C]. The culture media from RAW-ASC cells treated with nigericin showed increased levels of IL-1β and IL-18, compared to untreated cultures (p<0.0001; Fig. 8 [D-E]). However, adding hiPSC-NSC-EVs after nigericin treatment markedly attenuated IL-1β and IL-18 levels compared with the nigericin-only group (p<0.001-0.0001; Fig. 8 [D-E]). In contrast, adding hiPSC-NSC-EVs that are depleted of miR-30e-3p (knockdown EVs [KD-EVs]) did not reduce IL-1β and IL-18 levels compared to the nigericin alone group (p>0.05, Fig. 8 [D-E]). Similar results were observed for the measurement of IL-1β and IL-18 in the cell lysates of these cultures (Fig. 8 [F-G]).

**Figure 8:**
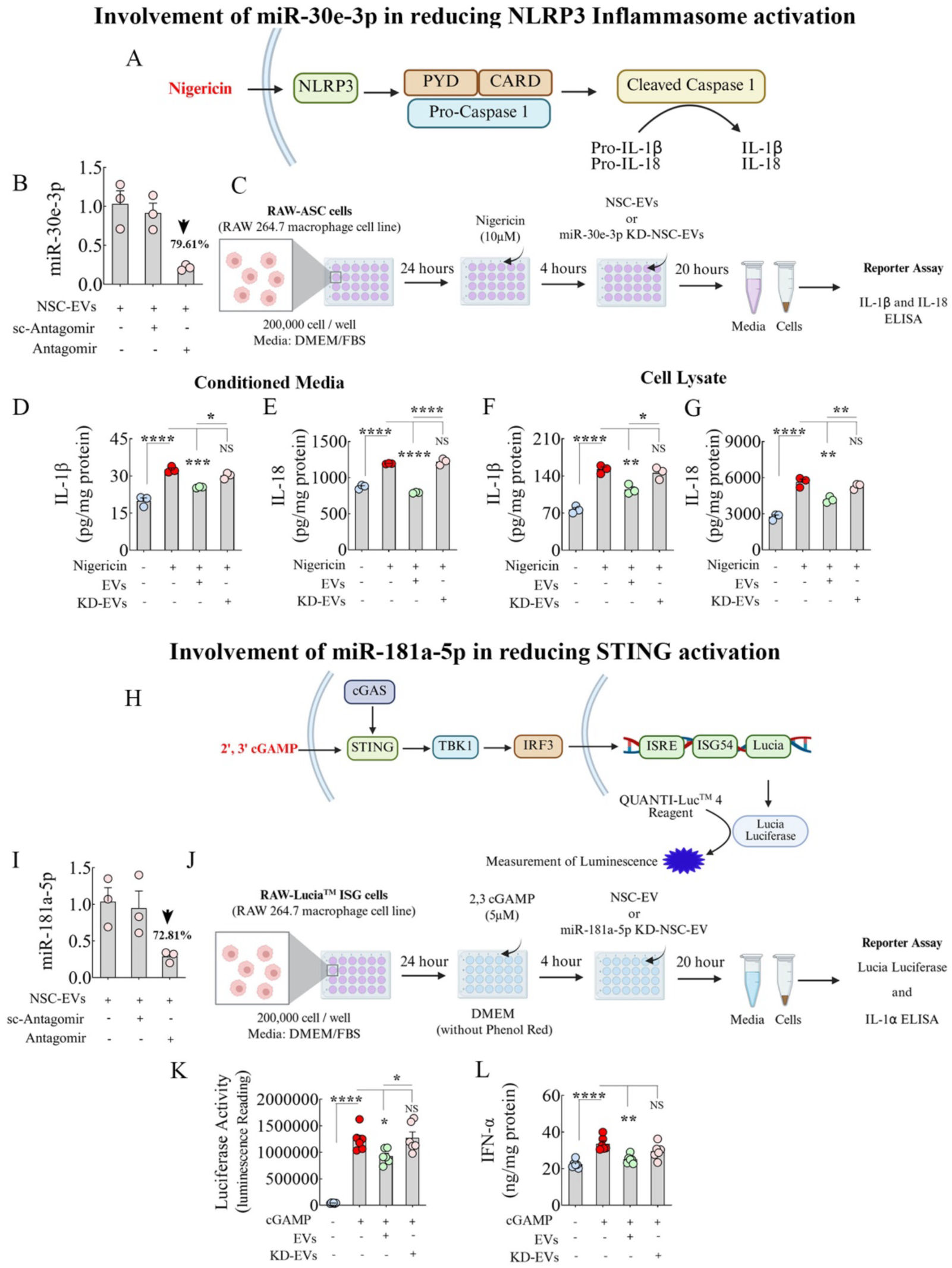
Depletion of miR-30e-3p in extracellular vesicles from human induced pluripotent stem cell-derived neural stem cells (hiPSC-NSC-EVs) diminished their ability to inhibit NLRP3 Inflammasome activation, and depletion of miR-181a-5p within hiPSC-NSC-EVs reduced their ability to inhibit STING activation. Cartoon A depicts the sequential changes following NLRP3 stimulation by Nigericin in RAW-ASC cells, culminating in the release of IL-1β and IL-18. Bar chart B compares the levels of miR-30e-3p in EVs derived from naive, miR-30e-3p antagomir-treated, or miR-30e-3p scrambled-antagomir-treated hiPSC-NSCs. Cartoon C illustrates the research design used to test the effects of naive hiPSC-NSC-EVs, hiPSC-NSC-EVs depleted of miR-30e-3p (KD-EVs), on RAW-ASC cells following nigericin stimulation. Bar charts D-G compare the concentrations of IL-1β (D, F) and IL-18 (E, G) in conditioned media (D-E), and cell lysates (F-G) of RAW-ASC cell cultures between various treatment groups. Cartoon H depicts the sequential changes following STING activation by 2’, 3’ cGAMP stimulation in RAW-Lucia-ISG cells, culminating in increased luciferase activity, implying the activation of the TBK-1-IRF3 axis and the production of type 1 IFNs. Bar chart I compares the levels of miR-181a-5p in EVs derived from naive, miR-181a-5p antagomir-treated, or miR-181a-5p scrambled-treated hiPSC-NSCs. Cartoon J illustrates the research design used to test the effects of naive hiPSC-NSC-EVs, hiPSC-NSC-EVs depleted of miR-181a-5p (KD-EVs), on RAW-Lucia-ISG cells following 2’, 3’ cGAMP stimulation. Bar chart K compares luciferase activity in conditioned media of RAW-Lucia-ISG cells between various treatment groups. Bar chart L compares IFN-α concentration in cell lysates of RAW-Lucia-ISG cells between various treatment groups.

Collectively, these findings suggested that miR-30e-3p is the primary active component in hiPSC-NSC-EVs that inhibit NLRP3 inflammasome activation.

### 3.12. Depletion of miR-181a-5p within hiPSC-NSC-EVs diminished their ability to inhibit STING activation

By employing RAW-Lucia-ISG cells, we investigated how hiPSC-NSC-EVs inhibit 2’,3’ cGAMP - induced STING activation (Fig. 8 [H]). For this, we first confirmed significantly reduced miR-181a-5p levels in EVs isolated from hiPSC-NSCs transfected with antagomirs targeting miR-181a-5p. EVs from transfected hiPSC-NSCs displayed a ∼72.81% decline in miR-181a-5p expression compared to EVs from naïve hiPSC-NSCs (Fig. 8 [I]). No such reduction was observed in EVs isolated from hiPSC-NSCs transfected with the scrambled miR-181a-5p antagomir. A schematic representation of the experimental steps is shown in Fig. 8 [J]. The culture media of RAW-Lucia-ISG cells treated with 2’,3’ cGAMP showed increased Lucia luciferase activity compared to untreated cultures (p<0.0001, Fig. 8 [K]). However, adding hiPSC-NSC-EVs after 2’,3’ cGAMP treatment markedly reduced Lucia luciferase activity compared with the 2’,3’ cGAMP-only group (p<0.05; Fig. 8 [K]). In contrast, adding hiPSC-NSC-EVs that are depleted of miR-181a-5p (KD-EVs) did not reduce Lucia luciferase activity compared to the 2’,3’ cGAMP-only group (p>0.05, Fig. [K]). We also measured IFN-α levels in lysates from RAW-Lucia-ISG cells across different groups. A significant increase in IFN-α concentration was observed in RAW-Lucia-ISG cells treated with 2’,3’ cGAMP alone compared to untreated cultures (p<0.0001, Figure 8 [L]). However, RAW-Lucia-ISG cells treated with 2’,3’ cGAMP and naïve hiPSC-NSC-EVs displayed a reduced concentration of IFN-α (p<0.01, Figure 8 [L]). In contrast, RAW-Lucia-ISG cells treated with 2’,3’ cGAMP and hiPSC-NSC-EVs that are depleted of miR-181a-5p did not show reduced IFN-α (p>0.05, Figure 8 [L]). Overall, these findings suggested that miR-181a-5p is the primary active component in hiPSC-NSC-EVs that inhibit STING activation.

### 3.13. Interaction with hiPSC-NSC-EVS triggered widespread transcriptomic changes in aged microglia

From sc-RNA-seq data, we first examined the t-SNE plots, which did not show clear separation between aged-Veh and aged-EVs groups (Suppl. Fig. 6). Analysis of DEGs (p<0.01) revealed upregulation of 896 genes and downregulation of 2025 genes in the aged-EVs group vis-à-vis the aged-Veh group (Fig. 9 [A]). The KEGG enrichment analysis (KEGG mouse 2019) of DEGs with p<0.01 between the aged-Veh and aged-EVs groups determined that these genes were mapped in 71 pathways, including Alzheimer’s disease, Oxidative phosphorylation, Toll-like receptor (TLR) signaling pathway, MAPK signaling pathway, TNF signaling pathway, C-type lectin receptor signaling pathway, mechanistic target of rapamycin (mTOR) signaling pathway, ForkHead box (FoxO) signaling pathway, Notch signaling pathway, Ras-related protein 1 (Rap1) signaling pathway, and Ubiquitin-mediated proteolysis (Fig. 9 [B]). Analysis of DEGs mapped in these pathways between the aged-Veh and aged-EVs groups revealed upregulation of numerous genes in Alzheimer’s Disease, in which the majority of DEGs were associated with mitochondrial function. Interestingly, all DEGs associated with oxidative phosphorylation were also significantly upregulated in the aged-EVs group compared to the aged-Veh group, implying enhanced mitochondrial activity following EVs treatment in late middle-aged mice (Fig. 9 [C-D]). Furthermore, hiPSC-NSC-EVs treatment in aged mice modulated the transcriptome of multiple neuroinflammatory pathways in microglia, where many DEGs in the TLR signaling pathway, MAPK signaling pathway, TNF signaling pathway, and the C-type lectin receptor signaling pathway were significantly downregulated in the aged-EVs group, implying reduced neuroinflammatory signaling with hiPSC-NSC-EVs treatment (Fig. 10 [A-D]). Furthermore, multiple DEGs in longevity-related pathways, including the mTOR signaling pathway, FoxO signaling pathway, and Notch signaling pathways, were significantly downregulated (p<0.01) in the microglia of the aged-EVs group (Fig. 10 [E-G]). Additionally, multiple DEGs in pathways such as Rap1 signaling and ubiquitin-mediated proteolysis were significantly downregulated following hiPSC-NSC-EVs treatment in late middle-aged mice (Suppl. Fig. 7 [A-B]).

**Figure 9:**
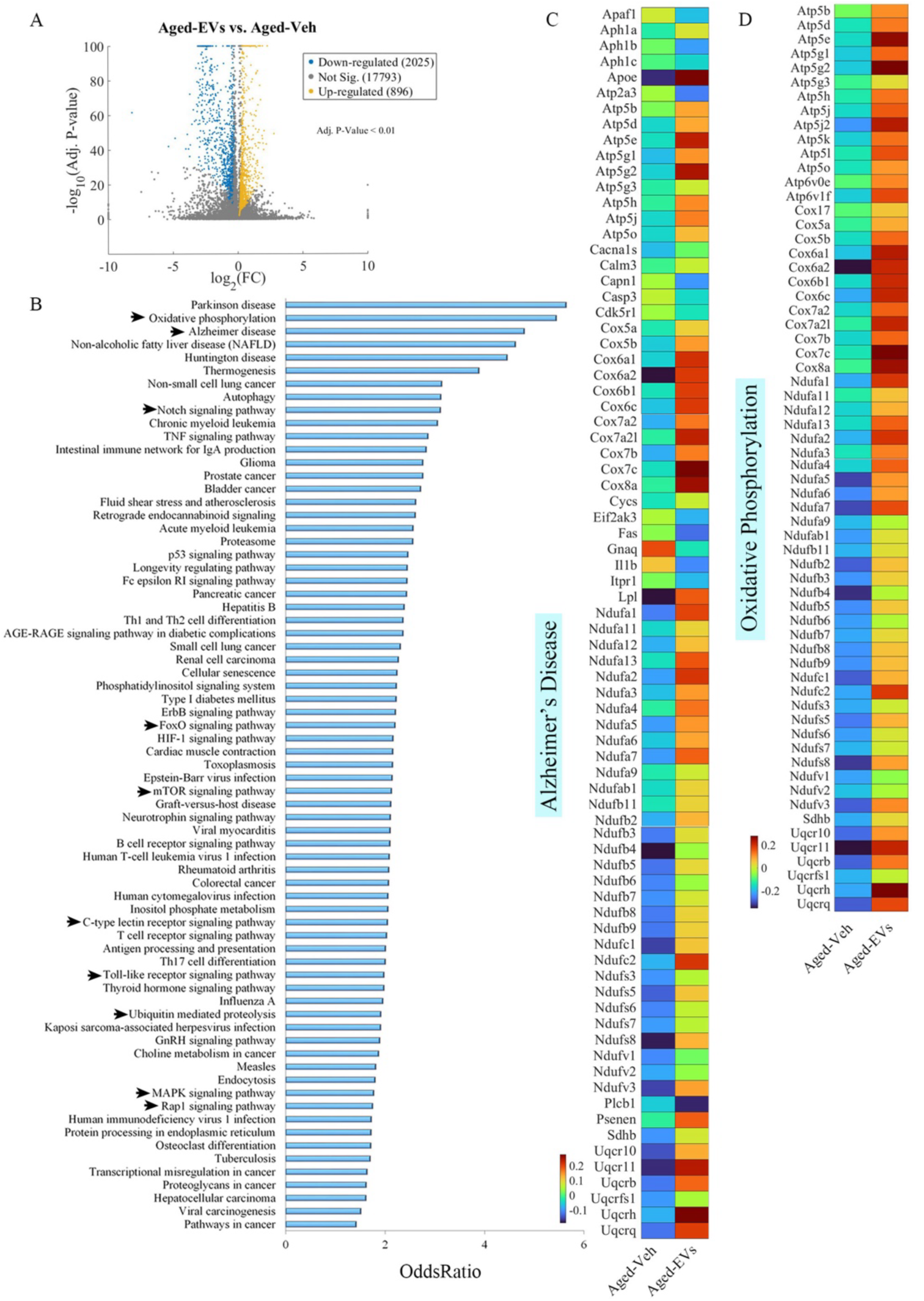
Intranasal administration of extracellular vesicles from human induced pluripotent stem cell-derived neural stem cells (hiPSC-NSC-EVs) induced widespread transcriptomic changes in microglia of late middle-aged male mice. The volcano plot in A displays differentially expressed genes (DEGs) that are significantly downregulated (blue) or upregulated (yellow) in the mouse from the Aged-EVs group compared to the mouse from the Aged-Veh group. Figure B illustrates the involvement of differentially expressed genes (DEGs) in the most significant pathways identified from KEGG enrichment pathway analysis. Heatmaps C and D compare the expression of various genes involved in pathways linked to Alzheimer’s disease (C) and oxidative phosphorylation (D) between the Aged-Veh and Aged-EVs groups.

**Figure 10:**
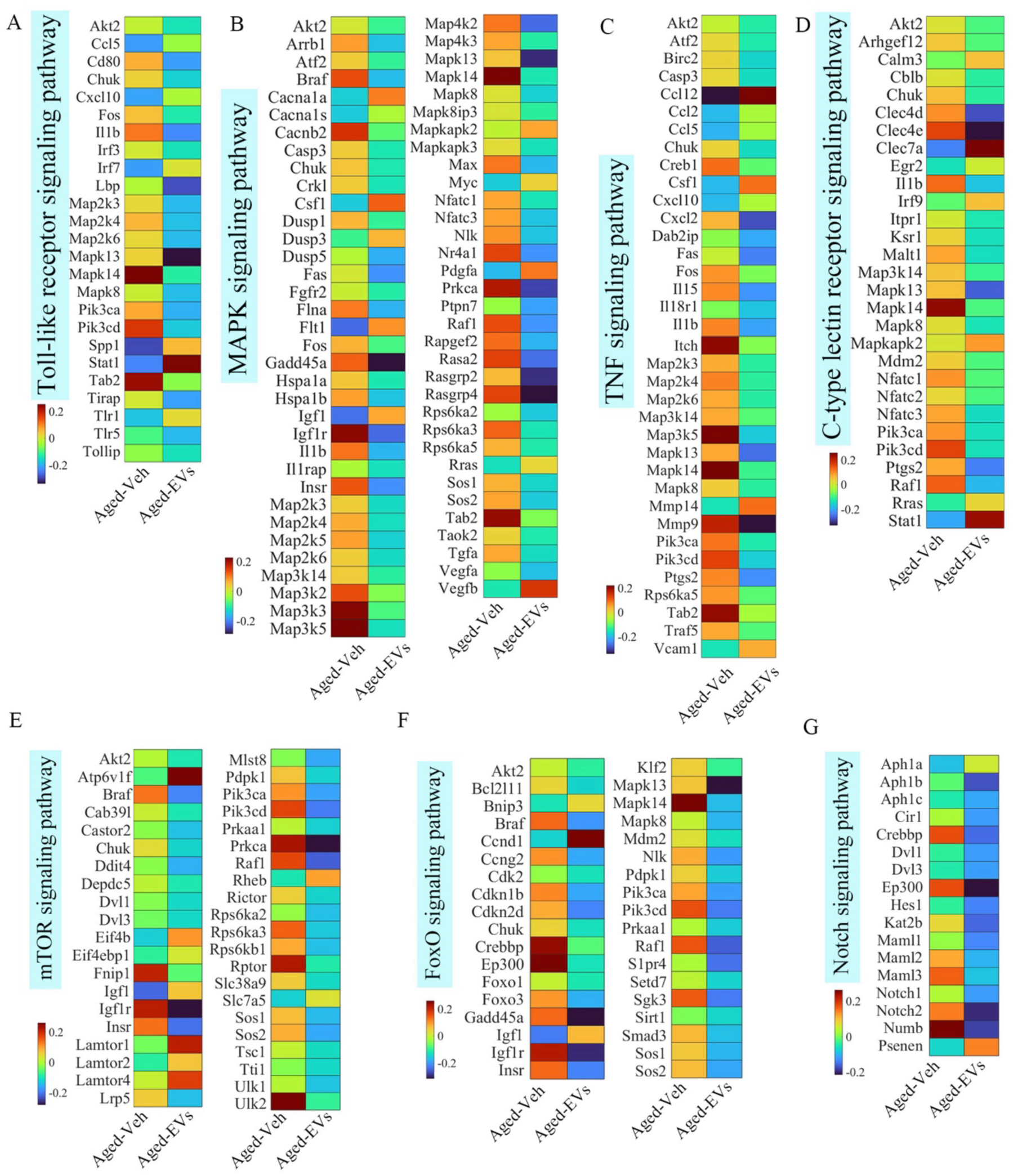
Intranasal administration of extracellular vesicles from human induced pluripotent stem cell-derived neural stem cells (hiPSC-NSC-EVs) reduced the expression of numerous genes implicated in many proinflammatory signaling pathways. Heatmaps in A-G compare the expression of genes involved in the Toll-like receptor (A), mitogen-activated protein kinase (MAPK) (B), tumor necrosis factor (TNF) (C), C-type lectin receptor (D), mechanistic target of rapamycin (mTOR) (E), ForkHead box (FoxO) (F), and Notch (G) signaling pathways between Aged-Veh and Aged-EVs groups.

We also performed microglial cluster extraction based on the DEGs between the clusters (Suppl. Table 3-6), which yielded five different clusters (Fig. 11 [A]). The KEGG pathway analysis of DEGs in clusters 1-5 revealed involvement in multiple signaling pathways, including those related to neuroinflammation (Suppl. Figs. 8-9). Among these, DEGs in cluster 1 are enriched for the TLR signaling pathway. The expression of genes such as *Pik3cd* and *Map3k8* was significantly reduced by hiPSC-NSC-EVs treatment in this microglial cluster (Fig. 11 [B]). The DEGs of cluster 2 are involved in IL-17 signaling and cholesterol metabolism. The expression of genes involved in IL-17 signaling, such as *Lcn2, S100a8,* and *S100a9*, was significantly reduced in microglia from the Aged-EVs group compared to microglia from the Aged-Veh group. Similarly, the expression of genes such as *Ldlr* and *Lpl,* involved in cholesterol metabolism, was significantly increased in microglia from the Aged-EVs group (Fig. 11 [C-D]). The DEGs of cluster 4 are found to be involved in IL-17 signaling, nucleotide-binding oligomerization domain (NOD)-like receptor signaling, and Rap1 signaling (Fig. 11 [E-G]). In IL-17 signaling, the expression of genes such as *Lcn2, S100a8,* and *S100a9* was significantly decreased in microglia from the Aged-EVs group (Fig. 11 [E]). In the NOD-like receptor pathway, the expression of genes such as *Gbp2, Oas1a,* and *Oas1g* was increased, and *Camp* was decreased in microglia from the Aged-EVs group (Fig. 11 [F]). In the Rap1 signaling, *Adcy8* and *Thbs1* genes were downregulated, and *Flt1* and *Igf1* were upregulated in microglia from the Aged-EVs group (Fig. 11 [G]). Thus, cluster analysis also revealed modulation of proinflammatory and cholesterol metabolism signaling pathways in microglia from mice receiving hiPSC-NSC-EVs.

**Figure 11:**
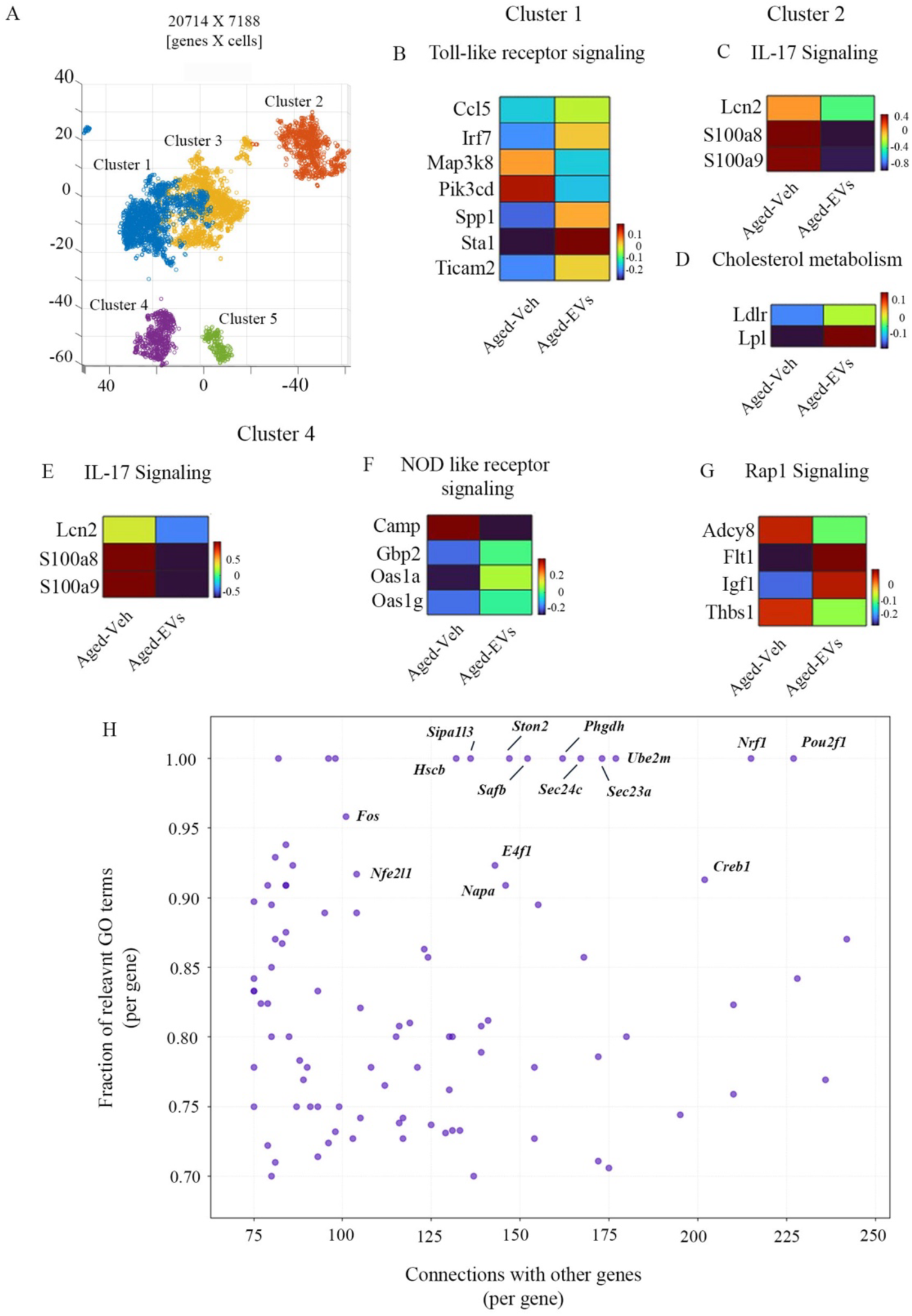
Figure A shows a t-SNE plot showing five distinct microglia clusters from late middle-aged male mice treated with extracellular vesicles from human induced pluripotent stem cell-derived neural stem cells (hiPSC-NSC-EVs) or vehicle. Heat maps in B-H compare the expression of genes identified in the most significant pathways identified from KEGG enrichment pathway analysis. These include the Toll-like receptor signaling pathway (B) from cluster 1; IL-17 (C) and cholesterol metabolism (D) signaling pathways from cluster 2; and IL-17 (E), nucleotide-binding oligomerization domain (NOD)-like receptor (F), and Ras-related protein 1 (Rap1) (G) signaling pathways from cluster 4. The scatter plot in H shows regulatory genes, identified through GeneWalk analysis, with various fractions of relevant gene ontology (GO) terms and variable connections to other genes.

Next, we performed GeneWalk analysis, yielding a scatterplot based on gene connections and fractions of relevant GO terms (Fig. 11 [H]). The most significant genes with >100 gene connections and the highest fractions of relevant GO terms (1:00) were identified. Among these, *Nrf1*, *Sec24C,* and *Fos* was downregulated in microglia from the Aged-EVs group compared to microglia from the Aged-Veh group (Fig. 11 [H]). These genes play significant roles in promoting microglia-mediated inflammatory pathways; hence, reduced expression of these genes is consistent with the diminished neuroinflammaging observed in the Aged-EVs group. In addition, the expression of the *Nfe2l1* gene was upregulated in the Aged-EVs group, which is significant because Nfe2l1 is known to maintain redox balance by stimulating antioxidant genes and maintaining mitochondrial homeostasis. Thus, GeneWalk analysis also confirmed a diminished proinflammatory transcriptome in microglia following treatment with hiPSC-NSC-EVs.

## 4. DISCUSSION

The results provide the first evidence that IN administration of hiPSC-NSC-EVs in late middle-aged mice can diminish the proinflammatory transcriptome within microglia in the aged brain. Such a change correlated with reductions in oxidative stress, mitochondrial dysfunction, and neuroinflammaging, and improved cognitive and memory function observed in aged mice receiving hiPSC-NSC-EVs. Furthermore, the antiinflammatory effects of hiPSC-NSC-EVs were associated with decreased activation of key signaling pathways, including NLRP3, p38/MAPK, cGAS-STING, JAK-STAT, and IFN-1 in the aged hippocampus.

### 4.1. Effect of hiPSC-NSC-EVs treatment on oxidative stress in the aged hippocampus

Increased oxidative stress is a key alteration in brain aging [10], characterized by elevated ROS levels that lead to mitochondrial dysfunction and synaptic damage in neurons [51]. Such changes contribute to cognitive impairments and the progression of neurodegenerative diseases like AD. Mitochondria generate most free radicals during energy production, and the accumulation of ROS disrupts redox homeostasis, leading to genomic instability and the transcription of many proinflammatory genes [10, 52]. Moreover, increased ROS can damage mitochondrial DNA, further impairing neuronal function [10, 52–53]. Therefore, strategies to reduce oxidative stress and protect the mitochondrial electron transport chain are crucial for preventing cognitive decline in old age. In this context, the current study shows that hiPSC-NSC-EVs treatment effectively reduces oxidative stress in the aged hippocampus of late-middle-aged male and female mice. These were evidenced by lower levels of MDA and PCs, which indicate reduced lipid peroxidation and protein oxidation, alongside increased NRF-2 and SOD levels, key components of antioxidant defense. Additionally, gene expression for many mitochondrial respiratory chain proteins was normalized in treated mice. The neuroprotective protein hemopexin, enriched in the hiPSC-NSC-EVs [26], may contribute to this protective effect against oxidative stress, given its role in safeguarding neurons from heme and ROS [54]. Additionally, diminished proinflammatory microglial transcriptome induced by hiPSC-NSC-EVs has likely contributed to reduced oxidative stress, as persistent neuroinflammation can elevate oxidative stress in the neuronal microenvironment. Such changes in microglia in the hippocampus of mice receiving hiPSC-NSC-EVs were evident from lower expression of numerous proinflammatory genes, higher expression of many genes that enhance oxidative phosphorylation, and reduced activation of several microglia-mediated proinflammatory signaling pathways.

### 4.2. Impact of hiPSC-NSC-EVs on neuroinflammatory signaling cascades in the aged hippocampus

Age-related neuroinflammation, or neuroinflammaging, is a significant aspect of brain aging [55–56]. In C57BL/6 mice, the incidence of neuroinflammation increases with age, accompanied by notable changes in microglia, including increased proliferation, reactivity, and motility, as well as altered gene expression [57–59]. Aging microglia express proinflammatory markers such as IL-1β, TNF-α, and IL-6, resulting in a chronic inflammatory environment detrimental to neuronal function [60–61]. Key neuroinflammatory pathways, particularly the NLRP3 inflammasome, are activated in this context [12–13]. Triggered by DAMPs like ROS and cellular debris, the NLRP3-ASC inflammasome complex activates caspase-1, leading to the maturation and increased release of IL-1β and IL-18 [15]. These changes initiate downstream signaling primarily through the p38/MAPK pathway, leading to the sustained release of proinflammatory cytokines and perpetuating chronic neuroinflammation [16–17]. Activation of the cGAS-STING-IFN-1 signaling is another pathway that plays a key role in neuroinflammaging [14].

Triggered by sensing dsDNA or mtDNA, this pathway leads to increased production of type 1-IFN, which is crucial for antiviral defense and immune regulation [14, 17, 62]. These interferons then signal through the JAK-STAT pathway to promote the transcription of interferon-stimulated genes (ISGs), thereby driving neuroinflammation and altered immune responses [14, 18, 63].

Thus, moderate chronic neuroinflammation, driven by NLRP3-p38/MAPK and cGAS-STING-IFN-1 signaling in the brain, significantly contributes to cognitive decline with aging. Targeting these pathways may help maintain cognitive function in old age. The current study found that hiPSC-NSC-EVs treatment in late-middle-aged mice reduced chronic neuroinflammation in the hippocampus, as evidenced by decreased proinflammatory transcriptomic signatures in microglia and a less proinflammatory microenvironment. First, treatment reduced microglial cluster formation and lowered the percentage of microglia with NLRP3 inflammasome complexes. Second, significant reductions in key proteins that activate the NLRP3 inflammasome and the p38/MAPK pathway were observed, including NLRP3, ASC, cleaved Caspase-1, pMAPK, and AP-1. Third, levels of proteins in the cGAS-STING-IFN-1 and JAK-STAT signaling cascades, such as p-cGAS, p-STING, p-IRF3, IFN-α, p-JAK1/2, and p-STAT1, as well as in ISG expression, were notably decreased. These changes were consistent in both male and female mice treated with hiPSC-NSC-EVs compared with vehicle-treated controls. Thus, hiPSC-NSC-EVs can slow chronic neuroinflammation in the aged brain by modulating the microglial proinflammatory transcriptome and inflammatory signaling cascades.

### 4.3. Direct effects of hiPSC-NSC-EVs on microglia in the aged hippocampus

Our earlier studies have shown that hiPSC-NSC-EVs can significantly lower IL-6 release from LPS-stimulated macrophages and IL-1β and TNF-α release from LPS-stimulated hiPSC-derived iMicroglia [26, 28]. Additionally, in neuroinflammatory conditions, such as in rodent models of status epilepticus or LPS-induced neuroinflammation, hiPSC-NSC-EVs markedly decreased microglia-mediated neuroinflammation [26, 29]. Our recent study also indicated that hiPSC-NSC-EVs can modulate disease-associated microglia in 5xFAD mice without affecting their phagocytic activity [17]. More importantly, in the current study, scRNA-seq analysis of microglia seven days after hiPSC-NSC-EVs administration showed that these EVs can alter the proinflammatory microglial transcriptome. hiPSC-NSC-EVs treatment enhanced the expression of genes related to AD, mitochondrial function, and cholesterol metabolism while reducing the expression of genes linked to neuroinflammatory pathways. Key pathways affected include AD and oxidative phosphorylation, with upregulation of genes crucial for mitochondrial respiratory chain integrity, which is beneficial as reduced expression of these genes is associated with mitochondrial dysfunction in aging and AD [64–66]. Increased expression of genes (*Ldlr and Lpl1*) related to cholesterol metabolism is also advantageous, as deficiency of these genes promotes microglial proliferation, activation, and lipid droplet accumulation [67–68]. The diminished proinflammatory pathways in microglia mediated by hiPSC-NSC-EVs include TLR, NOD, MAPK, TNF, C-type lectin receptor, and IL-17 signaling. Such effects have immense value, as aging, AD, or neurodegenerative diseases are associated with increased expression of genes promoting TLR [69], NOD [70], MAPK [71], TNF [72–73], and C-type lectin receptor [74] signaling pathways. Moreover, decreased expression of genes linked to NOD-signaling, such as Pik3cd and Map3k8, has therapeutic value, as they promote microglial activation [75–76]. Also, decreased expression of genes (Lcn2, S100a9, and S100a8) involved in IL-17 signaling is important, as these genes are implicated in the induction of a proinflammatory signature in microglia [77–79].

Furthermore, the expression of many genes associated with mTOR, FoxO, Notch, Rap1 signaling, and the ubiquitin-mediated proteolysis pathway was significantly downregulated in mice treated with hiPSC-NSC-EVs. These changes also carry significant implications for neuroinflammaging. For instance, heightened mTOR signaling in microglia contributes to a primed inflammatory state in the aged brain [80]. On the other hand, increased activity of FoxO transcription factors, such as FoxO3, can lead to greater microglial activation [81–82]. Conversely, elevated Notch signaling can further amplify microglial activation in neuroinflammatory conditions [83], and enhanced Rap1 signaling is known to drive neuroinflammation [84]. Interestingly, the reduced expression of genes related to the ubiquitin-mediated proteolysis pathway is unexpected, given its crucial role in clearing ubiquitylated protein aggregates [85]. This downregulation may reflect a compensatory mechanism or the brain’s effort to restore balance in a microenvironment characterized by lower oxidative stress and reduced neuroinflammation, mediated by hiPSC-NSC-EVs treatment. Additionally, GeneWalk analysis revealed four regulatory genes with over 100 connections and GO term scores exceeding 0.95, all relevant to microglial function. These include *Nrf1, Sec24c, Fos,* and *Nfe2l1*. Notably, the expression of *Nrf1, Sec24c,* and Fos was downregulated in aged mice receiving hiPSC-NSC-EVs, suggesting an advantage. First, decreased *Nrf1* expression in microglia can reduce the production of proinflammatory cytokines TNF-α and IL-1β [86]. Second, decreased *Sec24c* expression can reduce STING activation, as Sec24c is required for STING oligomerization [87]. Third, decreased *Fos* expression can reduce neuroinflammatory response in the hippocampus [88]. Furthermore, *Nfe2l1* expression was upregulated, which is significant because it helps maintain redox balance by stimulating antioxidant genes and supporting mitochondrial homeostasis [89]. Thus, hiPSC-NSC-EVs treatment in late middle-aged mice promoted robust antiinflammatory effects on microglia.

### 4.4. Mechanisms by which hiPSC-NSC-EVs alleviated neuroinflammation in the aged hippocampus

The miRNA and protein composition of hiPSC-NSC-EVs employed in this study provides insights into their potential antiinflammatory mechanisms in the aged hippocampus. Our previous studies have identified miRNAs, such as miR-21-5p and miR-103a, and proteins, such as pentraxin-3 (PTX3), hemopexin, and galectin-3 binding protein (Gal3BP), in hiPSC-NSC-EVs that can promote antiinflammatory effects [26,28]. For example, miR-21-5p can enhance antiinflammatory activity by modulating NF-κB signaling, increasing IL-10 levels, and inhibiting TNF-α release [90–93]. Whereas PTX3 can stimulate beneficial A2 astrocytes and help maintain the blood-brain barrier [94–96]. Additionally, miR-103a, hemopexin, and Gal3BP can substantially reduce neuroinflammation and promote the transition of microglia from a proinflammatory to an antiinflammatory state [22, 97–99]. More importantly, cell culture experiments with engineered RAW-ASC and RAW-Lucia-ISG cells in this study provide evidence that miR-30e-3p and miR-181a-5p are involved in the inhibition of the NLRP3 inflammasome and cGAS-STING pathway, respectively, by hiPSC-NSC-EVs. In RAW-ASC cells, naïve hiPSC-NSC-EVs reduced NLRP3 inflammasome activation and IL-1β and IL-18 release after nigericin stimulation, an effect lost with miR-30e-3p-depleted hiPSC-NSC-EVs. Similarly, in RAW-Lucia-ISG cells, naïve hiPSC-NSC-EVs decreased luciferase activity after cGAMP stimulation, indicating suppressed STING activation, which was absent when miR-181a-5p-depleted hiPSC-NSC-EVs were employed. Overall, multiple miRNAs and proteins within hiPSC-NSC-EVs can reduce microglial proinflammatory responses, with miR-30e-3p and miR-181a-5p specifically linked to inhibition of the NLRP3 inflammasome and the cGAS-STING pathway, respectively. These findings align with previous studies on the roles of these miRNAs [43–44].

### 4.5. Impact of hiPSC-NSC-EVs treatment in late middle age on cognitive function in old age

Cognitive and mood impairments are commonly observed in late-middle-aged mice [100–102]. At 18 months, both male and female mice exhibited deficits in recognition memory (measured by NORT) and object location memory (measured by OLT), which rely on the integrity of the perirhinal cortex and hippocampus [32,33]. These impairments persisted into old age for vehicle-treated mice. However, male and female mice treated with hiPSC-NSC-EVs showed improved recognition and object-location memory in old age, suggesting that hiPSC-NSC-EVs treatment induced beneficial changes in the hippocampus and other brain regions, thereby reversing cognitive deficits. Since increased oxidative stress and neuroinflammation are linked to cognitive impairments in aging mice [55, 103–106], the current study suggests that reductions in oxidative stress and neuroinflammatory signaling, mediated by hiPSC-NSC-EVs treatment, have improved cognitive function in old age. This finding aligns with the understanding that neuroinflammation is a risk factor for MCI, dementia, and AD [107]. However, any positive impact of hiPSC-NSC-EVs on neurons, which may also contribute to better cognitive function in aged mice, requires further investigation. Nonetheless, the current study is the first to demonstrate the beneficial effects of hiPSC-NSC-EVs on cognitive function in aged animals, whereas previous studies using EVs from mesenchymal stem cells have reported other benefits, such as improved locomotor function and neuroprotection [108] and microglia-mediated synapse remodeling [109] in aging models.

## 5. CONCLUSION AND LIMITATIONS

The results indicate that intranasal (IN) administration of hiPSC-NSC-EVs during late middle age can alter the proinflammatory transcriptome within microglia in the aged brain, leading to reduced oxidative stress and neuroinflammaging, along with improved cognitive and memory function. The antiinflammatory effects were demonstrated by decreased activation of several microglia-mediated proinflammatory signaling pathways in the aged hippocampus. Therefore, hiPSC-NSC-EVs treatment in late middle age has promise to help reduce brain inflammation and potentially delay MCI and AD. However, more research is needed before clinical application. Furthermore, although there was no interaction between sex and hiPSC-NSC-EVs treatment, minor sex-specific effects were observed, with female mice receiving hiPSC-NSC-EVs showing increased expression of a gene (Ndufs6) and higher concentration of a few proteins (CAT, pIRF3), compared with males, suggesting a need for further investigation into sex-based responsiveness. Future studies should also examine the effects of treatment initiation timing, various doses, and long-term efficacy of hiPSC-NSC-EVs. Additionally, the development of good manufacturing practice protocols for large-scale production and rigorous testing of clinical-grade hiPSC-NSC-EVs in larger animal models is essential for the successful translation of hiPSC-NSC-EVs therapy to older adults or those with MCI.

## ACKNOWLEDGEMENTS

The TEM images of hiPSC-NSC-EVs were taken at the Image Analysis Laboratory, Texas A&M Veterinary Medicine & Biomedical Sciences (RRIS: SCR_0222479).

## DECLARATION OF GENERATIVE AI AND AI-ASSISTED TECHNOLOGIES IN THE WRITING PROCESS

During the preparation of this manuscript, the authors utilized the Grammarly software primarily to check Grammar and spelling but did not use AI-assisted technologies to generate manuscript text. The authors take full responsibility for the content of the publication.

## AUTHOR CONTRIBUTIONS

Concept: AKS. Research design: AKS, LNM, MK, SR, SA, and RU. Data collection, analysis, and interpretation: LNM, MK, SR, SA, RU, GS, BS, YS, SVG, VSK, JEJ, PAS, AL, XR, JJC, and AKS. Preparation of figure composites: LNM, MK, SR, SA, and YS. Manuscript writing: LNM, MK, and AKS. All authors provided feedback, edits, and additions to the manuscript text and approved the final version of the manuscript.

## ETHICAL APPROVAL

The animal care and experimental procedures were conducted per the animal protocol approved by the Animal Care and the Use Committee (IACUC) of Texas A&M University School of Medicine.

## DATA AVAILABILITY

Source data are included in this original research article. Any additional data requests are available from the corresponding author upon request.

