## Supplemental File for "Intranasal Human NSC-Derived EVs Therapy Can Restrain Inflammatory Microglial Transcriptome, and NLRP3 and cGAS-STING Signaling, in Aged Hippocampus"

**CONTENTS**

1. Additional details to “Materials and Methods” section

2. Additional details to “RESULTS” section

3. References Cited

### 1. Additional information to "Materials and Methods" section:

#### *1.1. Human induced pluripotent stem cell (hiPSC)-derived neural stem cell (NSC) cultures*

The colonies derived from a human induced pluripotent stem cell (hiPSC) line, the IMR90-4 line obtained from the Wisconsin International Stem Cell Bank (Madison, WI, USA), were cultured for 24 hours. Subsequently, the medium was replaced with neural induction medium to differentiate the hiPSCs into neural stem cells (NSCs). The hiPSC line (i.e., the IMR90-4 line) was generated from fibroblasts obtained from a female donor. The NSCs were expanded and passaged every 7 days; those from different passages were cryoprotected and stored in liquid nitrogen. Passage 11 (P11) NSCs were cultured to 70% confluency, at which point they were seeded into 150 × 20 mm culture plates containing NSC expansion medium. Once the cultures achieved approximately 90% confluency, the spent media were harvested and stored at −20°C for extracellular vesicle (EVs) isolation. The purity of P11 NSCs was confirmed using immunofluorescence staining for NSC-specific markers, including nestin (anti-nestin, 1:1000; EMD Millipore, Burlington, MA, USA) and Sox-2 (anti-Sox-2, 1:300; Millipore).

#### *1.2. Isolation of extracellular vesicles (EVs) from hiPSC-NSC culture media using anion-exchange and size-exclusion chromatographic methods*

The EVs shed by hiPSC-NSCs were isolated using a series of protocols, including centrifugation, filtration, anion exchange chromatography (AEC), and size-exclusion chromatography (SEC) (Upadhya et al., 2020). First, the conditioned media (CM) from P11 hiPSC-NSC cultures were centrifuged at 3000 rpm for 10 minutes. The supernatants were next filtered through a 0.22 µm filter (GE Healthcare, Uppsala, Sweden) to eliminate cellular debris. Next, an AEC column (1.5 x 12 cm; Bio-Rad, CA, USA) was prepared by filling it with 10 mL of Q-Sepharose Fast Flow (GE Healthcare) and equilibrating with 100 mL of equilibration buffer. The concentrated CM was added, followed by a washing step. The EVs were selectively eluted with an elution buffer containing 50 mM Tris and 1,000 mM sodium chloride (NaCl) at pH 8.0. Fractions from the AEC were collected at a flow rate of 1 mL/min, and the elution of EVs was continuously monitored using nanoparticle tracking analysis (NTA). The fractions containing EVs were pooled, concentrated using an ultrafiltration device with a 50 kDa cut-off, and stored for further use. Following AEC, size-exclusion chromatography (SEC) was performed using a column that contained 20 mL of Sephacryl S-500 High Resolution (GE Healthcare). The EVs were size fractionated with a mobile phase composed of 50 mM phosphate buffer and 200 mM NaCl at pH 7.4. Fractions were collected at a flow rate of 1 mL/min. The elution of EVs was continuously monitored by quantifying total protein using the bicinchoninic acid (BCA) method and through NTA. The SEC fractions with the highest EV concentrations and the lowest protein content were pooled, concentrated, and stored at -20$^0$ C for future use.

#### *1.3. Characterization of hiPSC-NSC-EVs for EVs-specific markers and ultrastructure*

Our earlier studies provide detailed characterization of the antiinflammatory, antioxidant, neuroprotective, and neurogenic properties of hiPSC-NSC-EVs, along with their miRNA and protein composition (Upadhya et al., 2020, 2022; Rao et al., 2025). For hiPSC-NSC-EVs employed in this study, we examined EV-specific markers via Western blotting, which included the apoptosis-linked 2 interacting protein X (ALIX), CD81, and CD63, and a negative marker of EVs (i.e., a deep cellular protein marker calnexin) using methods described elsewhere (Upadhya et al., 2020; Ayyubova et al., 2023). Briefly, an aliquot of EVs (100 µl) was mixed with 100 µl of a mammalian protein extraction reagent (ThermoFisher Scientific, Waltham, MA, USA) and lysed as described previously (Madhu et al., 2019). The total protein concentration in the lysate was measured using the Pierce BCA Protein Assay Kit (ThermoFisher Scientific), and 40 µg of protein were loaded and separated on 4–12% NuPAGE Bis-Tris Gels (ThermoFisher Scientific). Next, proteins were transferred onto nitrocellulose membranes using ThermoFisher Scientific's iBlot2 gel transfer technology. The membranes were processed for protein detection using antibodies against CD63 (1:1000, BD Biosciences, San Jose, California, CA, USA), CD81 (1:1000, BD Biosciences), ALIX (1:1000, Santa Cruz, Santa Cruz, Dallas, TX, USA), and calnexin (1:1000; ThermoFisher). The protein signals were then detected using an IBright 1500 (Invitrogen, Waltham, MA, USA) and a chemiluminescent substrate kit (ThermoFisher Scientific). The lysate from hiPSC-NSCs was also processed in parallel to confirm the absence of deep cellular protein calnexin in EV preparations (Ayyubova et al., 2023). We performed transmission electron microscopy (TEM) to visualize the morphology of isolated hiPSC-NSC-EVs, as described in our previous reports (Upadhya et al., 2020; Kodali et al., 2023; Ayyubova et al., 2023).

#### *1.4. Processing of brain tissues for immunohistochemistry*

The brain tissues were fixed with 4% paraformaldehyde via intracardiac perfusion, treated with various concentrations of sucrose, and processed for cryostat sectioning. Thirty-micrometer-thick coronal brain tissue sections were stored at −20°C in cryobuffer until further use, as detailed in our previous reports (Rao et al., 2008; Hattiangady et al., 2011; Long et al., 2017; Upadhya et al., 2019; Attaluri et al., 2022; Ayyubova et al., 2023; Kodali et al., 2023; Madhu et al., 2024). From each animal, serial sections (every 20th slice) were taken through the septotemporal axis of the hippocampus for immunohistochemical studies aimed at identifying glial fibrillary acidic protein (GFAP)- positive astrocytes and IBA-1-positive microglia. The primary antibodies used were goat anti-IBA-1 (1:1000, Abcam, Cambridge, MA, USA) and rabbit anti-GFAP (1:3000, Agilent Technologies, Carpinteria, CA, USA). For the secondary antibodies, we used horse biotinylated anti-rabbit or anti-goat IgG (1:250 dilution, Vector Labs, Burlingame, CA, USA). The avidin-biotin complex reagent and the chromogen Vector Gray were also obtained from Vector Labs. The brain tissue sections were mounted on gelatin-coated slides, air-dried, counterstained with nuclear fast red (Vector Labs), dehydrated, cleared, and coverslipped with Permount. Finally, the sections were examined under a Nikon E600 microscope (Ayyubova et al., 2023; Kodali et al., 2023).

#### *1.5. Processing of brain tissues for single, dual or triple immunofluorescence methods*

Single-, dual-, or triple-immunofluorescence procedures were utilized to visualize various markers. The specific markers included: 1) IBA-1+ microglia and GFAP+ astrocytes that incorporated or interacted with PKH26+ hiPSC-NSC-EVs, and 2) microglia exhibiting the nucleotide-binding domain leucine-rich repeat and pyrin domain-containing receptor 3 (NLRP3) in conjunction with apoptosis-associated speck-like protein containing a CARD (ASC) complex. The tissue sections were washed in phosphate-buffered saline (PBS), blocked with 10% normal donkey serum, and incubated overnight at 4°C with individual primary antibodies for single immunofluorescence or a cocktail of two or three primary antibodies for dual or triple immunofluorescence. Following this, the sections were incubated with corresponding secondary antibodies, rinsed in PBS, and coverslipped using a slow fade/antifade mounting medium (Invitrogen). The primary antibodies employed in this study included: anti-goat IBA-1 (1:1000, Abcam), anti-goat NLRP3 (1:500, Millipore), anti-mouse ASC (1:500, Santa Cruz), and anti-rabbit IBA-1 (1:1000, Abcam). The secondary antibodies used were: donkey anti-rabbit IgG tagged with Alexa Fluor 488 (1:200, Invitrogen), donkey anti-goat IgG conjugated with Alexa Flour 594 (1:200, Invitrogen), donkey anti-goat IgG tagged with Alexa Fluor 488 (1:200, Invitrogen), donkey anti-mouse IgG tagged with Alexa Flour 594 (1:200, Invitrogen), and donkey anti-rabbit IgG tagged with Alexa Flour 405 (1:200, Invitrogen).

***1.6. Processing of hippocampal tissues for biochemical assays and qPCR studies***

The hippocampus from each brain was micro-dissected, lysed via 15–20 seconds of sonication at 4°C in a tissue extraction reagent (Invitrogen) containing protease/phosphatase inhibitor (1:100 dilution, ThermoFisher Scientific), and centrifuged for 10 min at 15,000 g. The supernatant was aliquoted and stored at −80 °C until further use (Madhu et al., 2021, 2024; Kodali et al., 2023). The RNA isolation kits were used to isolate total RNA from hippocampal tissues (Qiagen, Germantown, MD, USA). The cDNAs were generated by converting samples of RNA (500ng/µL) using RT2 First Strand Kit (Qiagen). The qRT-PCR was performed using RT² SYBR Green qPCR Mastermix and Primer mix (GeneCopoeia, Rockville, Maryland, USA) to measure the expression of genes linked to mitochondrial respiratory chain *(Ndufs6, Ndufs7, Sdha, Sdhb, Cyc1, Bcs1l, Cox7b, Cox4i2, Slc25a1, Atp6ap1),* inflammasome markers (NLRP3, PYCARD, Casp-1, IL-1β, IL-18) (Madhu et al., 2024).

***1.7. Interrogation of object recognition memory using a novel object recognition test***

A detailed methodology for the novel object recognition test (NORT) can be found in our previous studies (Hattiangady et al., 2014; Long et al., 2017; Upadhya et al., 2019; Madhu et al., 2024). Animals from all groups (n = 12-18 per group) participated in three trials, designated Trials 1-3 (T1-T3). In T1, the animals explored an open-field apparatus for 5 minutes. T2 commenced 15 minutes after T1, during which two identical objects were placed diagonally in the open field, and the animals explored these objects for another 5 minutes (see Fig. 7 [A]). T3 began 15 minutes later, with the animals being placed in the center of the open field. In this trial, one of the objects from T2 remained in its original location, while the other was replaced with a new object (see Fig. 7 [A]). The behavior of each mouse during T2 and T3 was monitored using the AnyMaze video-tracking system. We recorded the time spent with the familiar object (FO) and the novel object (NO), as well as the total object exploration time (TOET) for both T2 and T3. The percentages of exploration times spent with the FO compared to the NO were analyzed statistically within each group. To ensure the validity of the test, which relies on the animals exploring the two identical objects in T2 (see Fig. 7 [A]), only those animals that explored the objects for a minimum of 20 seconds in T2 and 8 seconds in T3 were included in the data analysis. Most animals (10-16 per group) met these criteria.

***1.8. Investigation of object location memory using an object location test***

A detailed protocol for conducting an object location test (OLT) is described in our previous studies (Hattiangady et al., 2014; Long et al., 2017; Upadhya et al., 2019; Kodali et al., 2023; Madhu et al., 2024). In this experiment, animals (n=12-18 per group) underwent three trials (T1-T3). In Trial 1 (T1), the animals were placed in an open-field apparatus for 5 minutes without any objects. After a 15-minute break, they proceeded to Trial 2 (T2), during which they explored two identical objects placed in the open field for another 5 minutes (see Fig. 7 [J]). Following another 15-minute interval, Trial 3 (T3) began, where the animals were placed again in the center of the open-field apparatus. In T3, one object remained in its original location from T2, while the other object was moved to a new location (see Fig. 7 [J]). The behavior of each mouse during T2 and T3 was monitored using the AnyMaze video-tracking system. Various results were calculated, including time spent with the object in the familiar location (OIFP) compared to the novel location (OINP), as well as total object exploration time (TOET) for both T2 and T3. The percentages of TOETs spent with the OINP relative to the OIFP were then computed for each group. Since success in this task depends on the careful exploration of the two distinct objects during T2 (see Fig. 7 [J]), only animals that spent at least 20 seconds exploring the objects in T2 and 8 seconds in T3 were included in the data analysis. Most animals (10-17 per group) satisfied these criteria.

#### 1.9. Single-cell RNA-sequencing of Microglia, alignment and filtering

The reads of scRNA libraries were aligned to the mm10 reference genome using Cell Ranger (version 7.0). The Count matrices of the Aged-Veh and Aged-EVs groups were further processed using scGEAToolbox (Cai, 2019; Cai and Osorio, 2021). Enrichment analysis of the differentially expressed genes (DEGs) was performed using Enrichr software. Briefly, we first imported and preprocessed scRNA-seq data into a SingleCellExperiment (SCE) object or a raw expression matrix, ensuring quality control using functions like sc_qcfilter to remove low-quality cells and genes, and then selecting highly variable genes via sc_hvg to focus on informative features. Next, dimensionality reduction and clustering were performed using tools such as sc_tsne for embeddings and sc_clusters for grouping cells into clusters, providing the group labels necessary for DE comparisons, such as the aged-Veh and aged-EVs groups. Next we performed core DE analysis by invoking the sc_deg or sc_degenes function, which takes inputs including the expression matrix X (genes by cells), gene list g, and group identifiers c, applying the Wilcoxon rank-sum test (also known as Mann-Whitney U test) to non-parametrically compare expression distributions between defined groups, computing metrics like p-values, adjusted p-values for multiple testing (e.g., via Benjamini-Hochberg false discovery rate), mean expressions, and log2 fold changes. Following computation, results were filtered based on significance criteria, such as p-values less than 0.05 and fold changes greater than 1.0, to identify differentially expressed genes (DEGs), which can then be visualized in the scGEATOOL graphical user interface using volcano plots. Later, downstream analyses, such as pathway enrichment to interpret biological implications, were performed.

**1.10. Power analyses performed to determine the number of animals**

The number of animals per group for neurobehavioral studies in this study was determined via power analysis using data from our previous studies (effect size [delta] = 1.4; alpha = 0.05) G* Power software, which indicated that to achieve a power of 0.8, final data are needed from 10 mice/gender/group. We included final data from 10-17 mice/gender/group after accounting for attrition due to some animals not reaching the criteria (e.g., lack of exploration of objects for significant periods). In the current study, the average effect size was 1.24 for males and 1.03 for females, with animal numbers ranging from 10 to 12 for males and 13 to 17 for females. For brain tissue analysis using histological and biochemical measures, a power analysis based on data from our previous studies indicated that 6 mice/gender/group are needed to achieve a power of 0.8 (effect size = 1.8, alpha = 0.05). We collected data from 6 mice/gender/group in this study.

### 2. Additional information to "Results" section

**2.1. Supplement Figure 1**

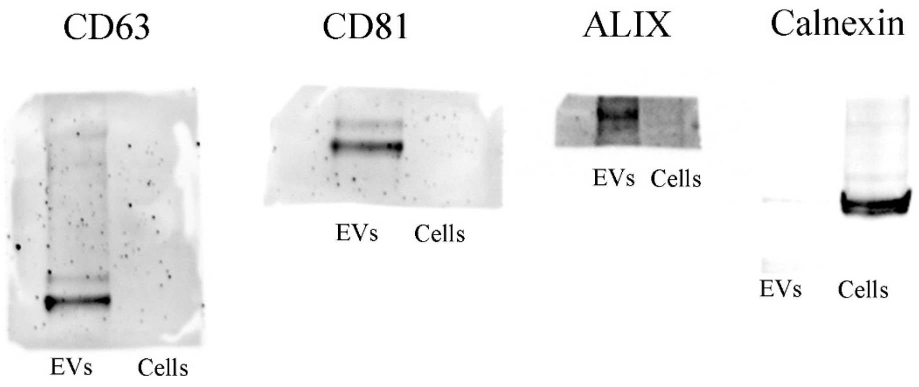

**Supplement Figure 1:** This figure displays the original uncropped western blot images of the EV marker proteins CD63, CD81, ALIX, and the EV negative marker calnexin.

### 2.2. Supplement Figure 2

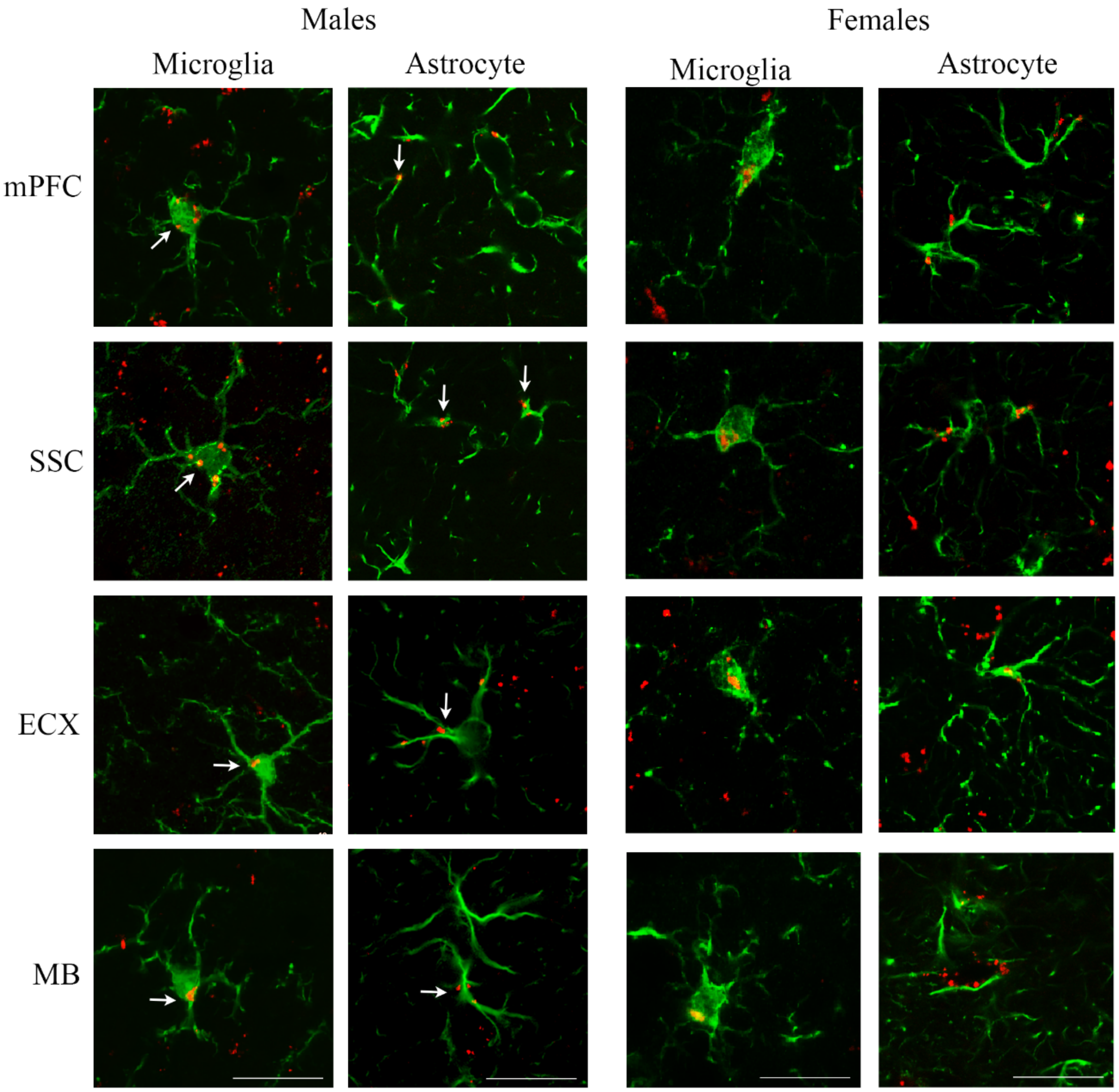

**Supplement Figure 2:** The images in the left two panels illustrate the incorporation/interaction of hiPSC-NSC-EVs into IBA-1+ microglia (panel 1) and GFAP+ astrocytes (panel 2) in the medial prefrontal cortex (mPFC), somatosensory cortex (SSC), entorhinal cortex (ECX), and midbrain (MB) of male mice. The images in the right two panels illustrate the incorporation/interaction of hiPSC-NSC-EVs into IBA-1+ microglia (panel 3) and GFAP+ astrocytes (panel 4) in mPFC, SSC, ECX, and MB of female mice. Scale bar = 10 µm.

### 2.3. Supplement Figure 3

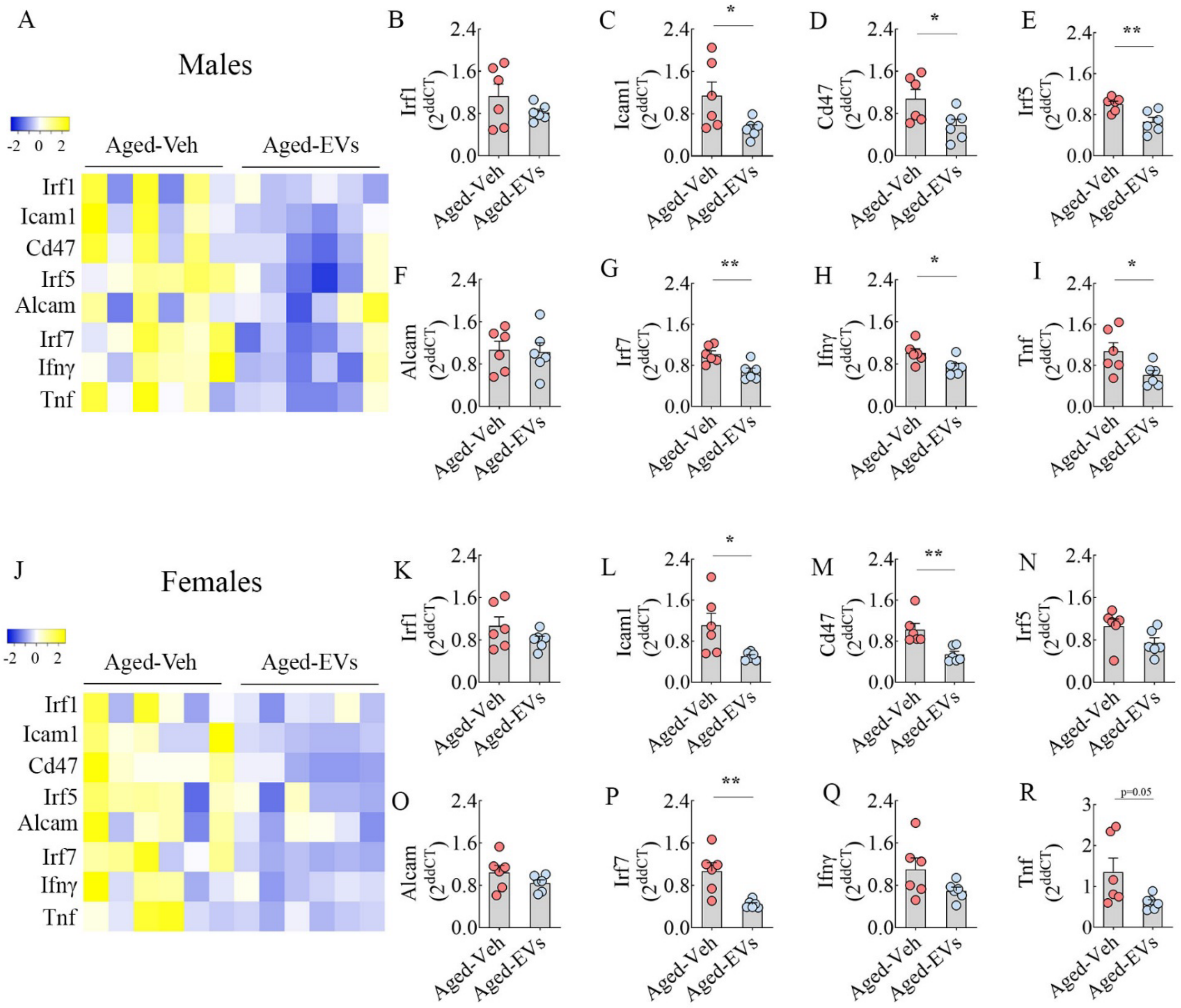

**Supplement Figure 3:** The heatmaps in A and J compare the relative expression of various interferon stimulatory genes (ISGs) between Aged-Veh and Aged-EVs groups. The bar charts (B-I, K-R) compare the relative expression of ISGs, including *Irf1, Icam1, Cd47, Irf5, Alcam, Irf7, Ifnγ,* and *Tnf,* in male (A-I) and female (J-R) mice between the Aged-Veh and Aged-EVs groups. *, p<0.05; and **, p<0.01.

### 2.4. Supplement Figure 4

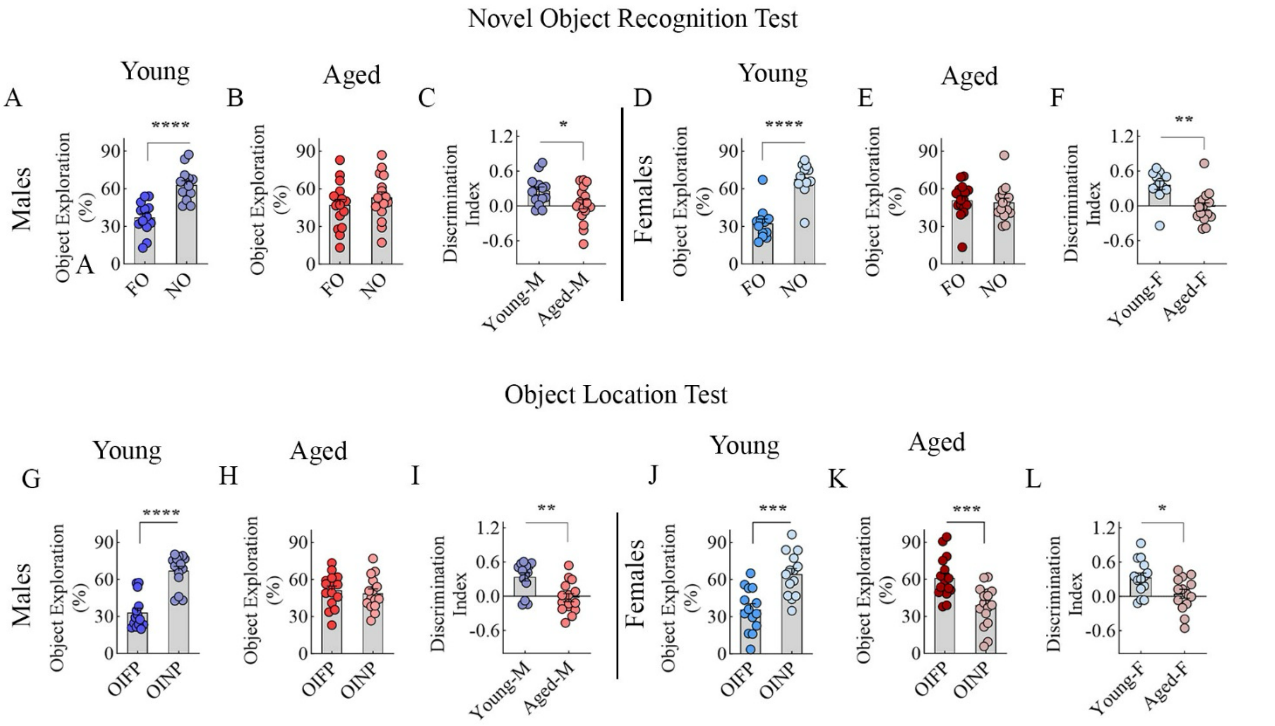

**Supplemental Figure 4:**

***Late middle-aged (18-month-old) male and female mice exhibited object recognition memory impairment compared to young adult male and female mice in a novel object recognition test (NORT).*** The bar charts A-B and D-E compare percentages of object exploration times spent with the familiar object (FO) vis-à-vis the novel object (NO) in young and late middle-aged males (A and B) and young and late middle-aged females (D and E). The bar charts C and F compare the NO discrimination index across young and late middle-aged males (C) and young and late middle-aged females (F).

***Late middle-aged (18-month-old) male and female mice exhibited impaired hippocampus-dependent cognitive function compared to young adult male and female mice in an object location test (OLT).*** The bar charts G-H and J-K compare the percentages of object exploration time spent with the object in the familiar place (OINP) versus the novel place (OINP) in young and late middle-aged males (G and H) and young and late middle-aged females (J and K). The bar charts I and L compare the OINP discrimination index across young and late middle-aged males (I) and young and late middle-aged females (L). *, p<0.05; **, p<0.01; ***, p<0.001; and ****, p<0.0001.

**2.5. Supplement Figure 5**

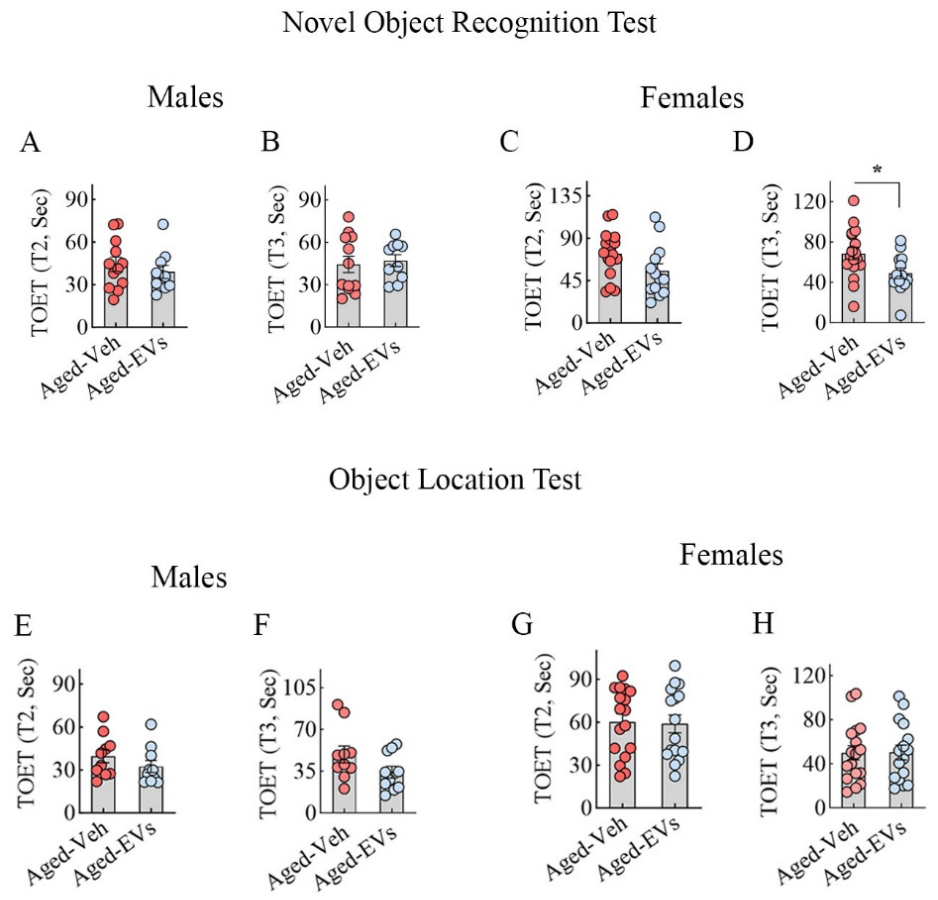

***Supplement Figure 5:*** The bar charts in A-D compare total object exploration times (TOETs) in T2 (A, C) and T3 (B, D) between Aged-Veh and Aged-EVs groups in males (A-B) and females (C-D) in a novel object recognition test (NORT). The bar charts in E-H compare total object exploration times (TOETs) in T2 (E, G) and T3 (F, H) between Aged-Veh and Aged-EVs groups in males (E-F) and females (G-H) in an object location test (OLT). *, $p<0.05$.

### 2.6. Supplement Figure 6

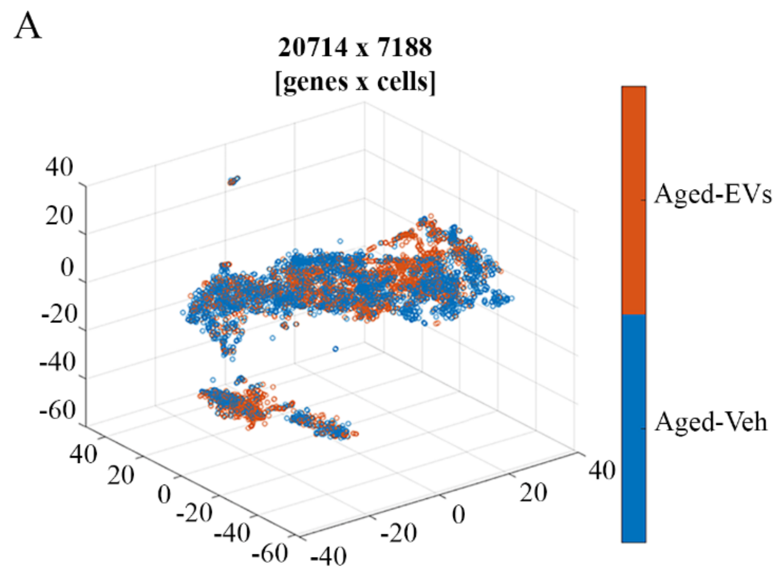

**Supplement Figure 6:** A t-SNE plot displaying the distribution of microglia in Aged-Veh and Aged-EVs groups from the single-cell RNA sequencing experiment.

**2.7. Supplement Figure 7**

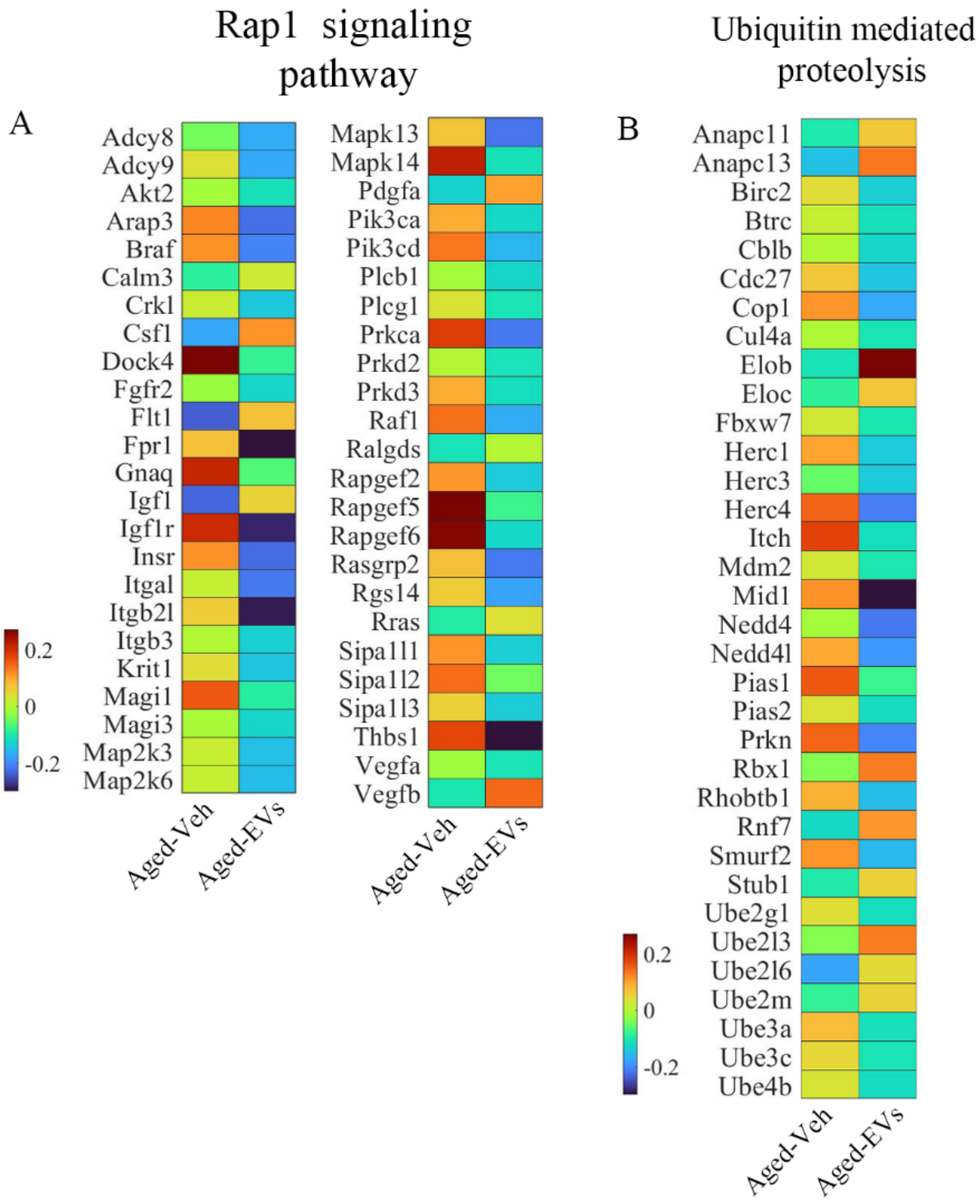

**Supplement Figure 7:** Heatmaps comparing the expression of various genes between Aged-Veh and Aged-EVs groups for Rap1 and the Ubiquitin-mediated proteolysis signaling pathways.

### 2.8. Supplement Figure 8

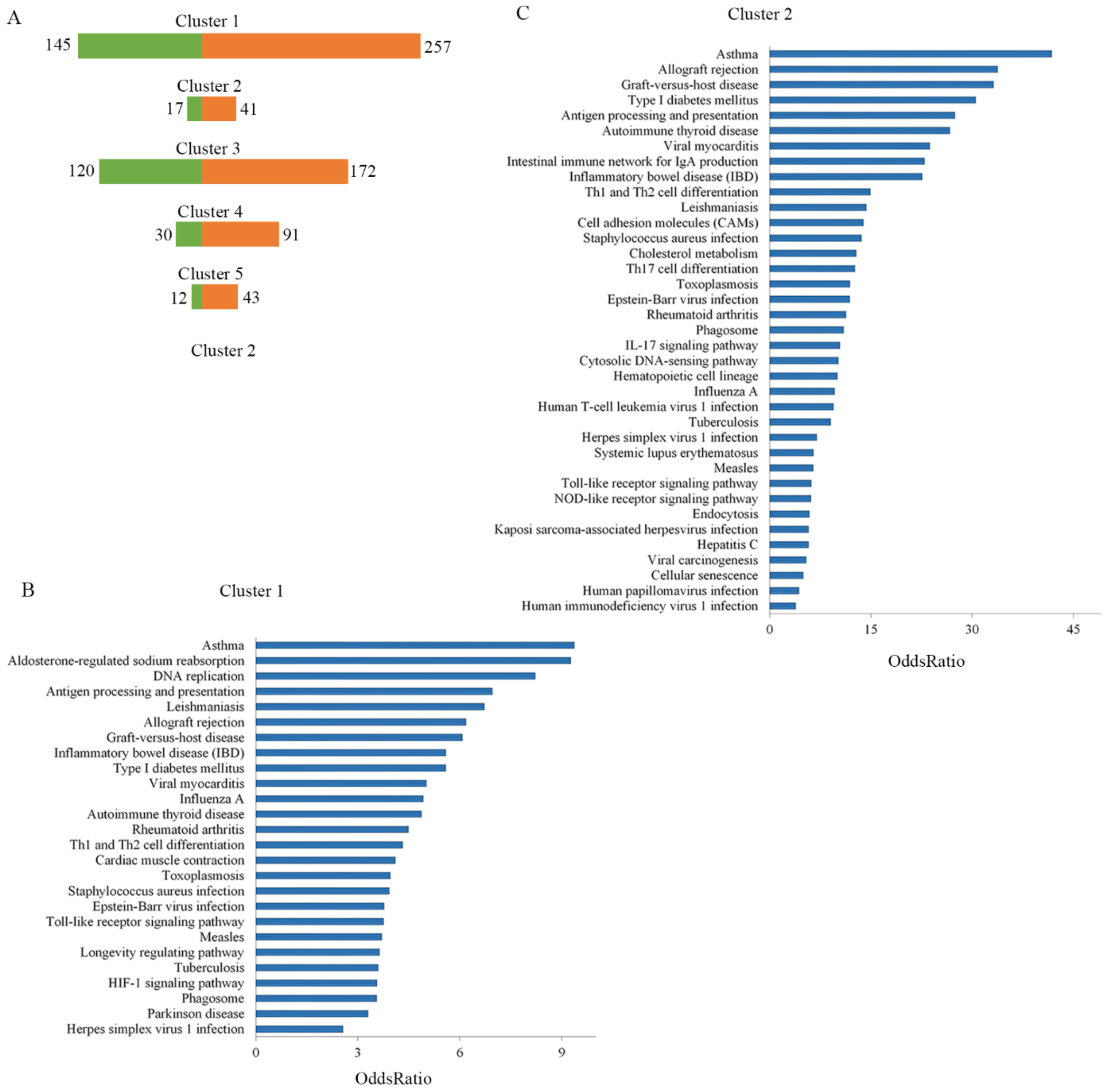

**Supplement Figure 8:** The bar graph in A shows the number of up-regulated and down-regulated genes in each microglial cluster. The bar graphs in B and C show the significant KEGG pathways identified in microglial clusters 1 and 2.

### 2.9. Supplement Figure 9

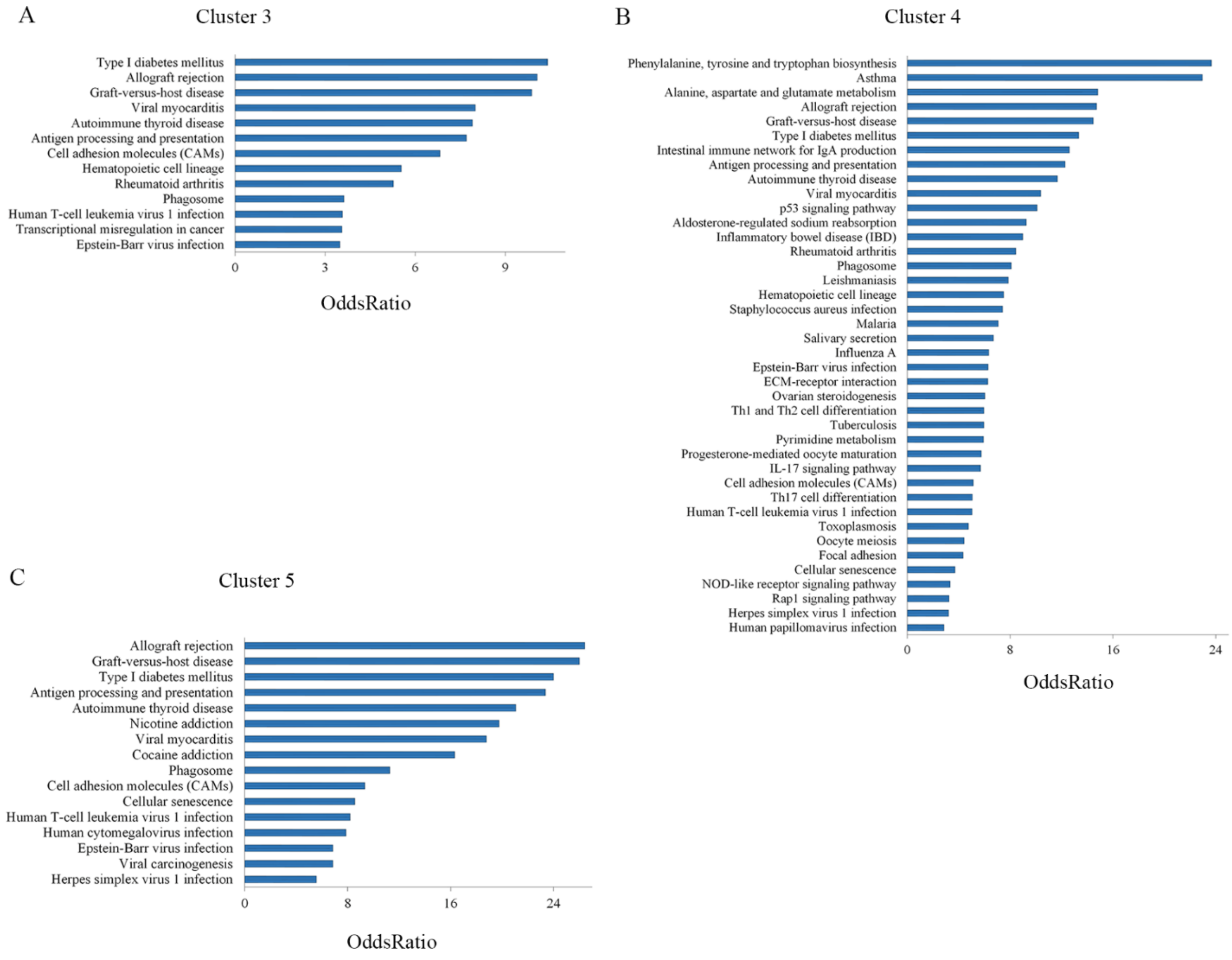

**Supplement Figure 9:** The bar graphs in A-C illustrate the significant KEGG pathways identified in microglial clusters 3,4, and 5.

### 2.10. Supplementary Table 1

#### List of genes mentioned in the manuscript

| Gene symbol | Gene Name |
|---|---|
| Adcy8 | Adenylate cyclase 8 |
| Alcam | Activated leukocyte cell adhesion molecule |
| Atp6ap1 | ATPase H+ transporting accessory protein 1 |
| Bcs1l | BCS1 homolog, ubiquinol-cytochrome C reductase complex chaperone |
| Casp1 | Caspase 1 |
| Cd47 | Cluster of differentiation 47 |
| Cox4i2 | Cytochrome C oxidase subunit 4i2 |
| Cox7b | Cytochrome c oxidase subunit 7B |
| Cyc1 | Cytochrome C1 |
| Flt1 | Fms-related receptor tyrosine kinase 1 |
| Fos | Transcription Factor AP-1 Subunit C-Fos |
| Gapdh | Glyceraldehyde-3-phosphate dehydrogenase |
| Gbp2 | Guanylate-binding protein 2 |
| Icam1 | Intercellular adhesion molecule 1 |
| Ifnγ | Interferon gamma |
| Igf1 | Insulin-like growth factor 1 |
| Il18 | Interleukin-18 |
| Il1β | Interleukin-1 beta |
| Irf1 | Interferon regulatory factor 1 |
| Irf5 | Interferon regulatory factor 5 |
| Irf7 | Interferon regulatory factor 7 |
| Lcn2 | Lipocalin 2 |
| Ldlr | Low-density lipoprotein receptor |
| Lpl | Lipoprotein lipase |
| Map3k8 | Mitogen-activated protein kinase kinase kinase 8 |
| Ndufs6 | NADH: ubiquinone oxidoreductase subunit S6 |
| Ndufs7 | NADH: ubiquinone oxidoreductase subunit S7 |
| Nfe2l1 | Nuclear factor erythroid 2-related factor 1 |
| Nlrp3 | Nucleotide-binding domain leucine-rich repeat (NLR) family pyrin domain-containing 3 |
| Nrf1 | Nuclear respiratory factor 1 |
| Oas1a | 2'-5' oligoadenylate synthetase 1A |
| Oas1g | 2'-5' oligoadenylate synthetase 1G |
| Pik3cd | Phosphatidylinositol-4,5-bisphosphate 3-kinase catalytic subunit delta |
| Pycard | PYD And CARD domain-containing |
| S100a8 | S100 Calcium binding protein A8 |
| S100a9 | S100 Calcium binding protein A9 |
| SdhA | Succinate dehydrogenase A |
| SdhB | Succinate dehydrogenase B |
| Sec24C | SEC24 Homolog C, COPII Component |
| Slc25a10 | Solute Carrier Family 25 Member 10 |
| Thbs1 | Thrombospondin 1 |
| Tnf | Tumor necrosis factor |

#### 2.11. Supplementary Table 2

**Sex-Specific Effects of hiPSC-NSC-EVs or Vehicle Treatment Analyzed via Two-way ANOVA**

| ***Comparisons*** | | Sex specific effects | Interaction between sex and hiPSC-NSC-EVs treatment |
|---|---|---|---|
| ***Astrocyte hypertrophy*** | Astrocyte area fraction (AF) | Yes (p<0.05) Greater in males in aged-Veh group | Yes (p<0.05) *Both males and females showed positive response* |
| ***Microglial numbers and clusters*** | Microglia count (Hippocampus) | No (p>0.05) | No (p>0.05) |
| | Microglia clusters (hippocampus) | No (p>0.05) | No (p>0.05) |
| ***Oxidative stress, NRF-2 and antioxidant markers*** | MDA | No (p>0.05) | No (p>0.05) |
| | PC | No (p>0.05) | No (p>0.05) |
| | NRF2 | No (p>0.05) | No (p>0.05) |
| | SOD | No (p>0.05) | No (p>0.05) |
| | CAT | Yes (p<0.01) *Greater in females in aged-EVs group* | Yes (p<0.01) *Only females showed positive response* |
| ***Expression of mitochondrial respiratory chain genes*** | Ndufs6 | Yes (p<0.05) Greater in females in aged-EVs group | Yes (p<0.05) *Both males and females showed positive response* |
| | Ndufs7 | No (p>0.05) | No (p>0.05) |
| | Sdha | No (p>0.05) | No (p>0.05) |
| | Sdhb | No (p>0.05) | No (p>0.05) |
| | Cyc1 | No (p>0.05) | No (p>0.05) |
| | Bcs1l | No (p>0.05) | No (p>0.05) |
| | Cox7b | No (p>0.05) | No (p>0.05) |
| | Cox4i2 | No (p>0.05) | No (p>0.05) |
| | Slc25a1 | No (p>0.05) | No (p>0.05) |
| | Atp6ap1 | No (p>0.05) | No (p>0.05) |
| ***Expression of NLRP3 inflammasome activation genes*** | Nlrp3 | No (p>0.05) | No (p>0.05) |
| | Pycard | No (p>0.05) | No (p>0.05) |
| | Caspase | No (p>0.05) | No (p>0.05) |
| | Il-18 | No (p>0.05) | No (p>0.05) |
| | Il-1β | No (p>0.05) | No (p>0.05) |
| ***Percentages of microglia displaying NLRP3 inflammasome complexes*** | Inflammasome in microglia | No (p>0.05) | No (p>0.05) |

| | | | |
|---|---|---|---|
| ***Mediators and end productions of NLRP3 inflammasome activation*** | NFkB-p65 | No (p>0.05) | No (p>0.05) |
| | NLRP3 | No (p>0.05) | Yes (p<0.05)<br>*Both males and females showed positive response* |
| | ASC | No (p>0.05) | No (p>0.05) |
| | Cleaved Caspase 1 | No (p>0.05) | No (p>0.05) |
| | IL-18 | No (p>0.05) | No (p>0.05) |
| | IL-1β | No (p>0.05) | No (p>0.05) |
| ***Mediators and end products of p38/MAPK signaling activation*** | Myd88 | No (p>0.05) | No (p>0.05) |
| | Ras | No (p>0.05) | No (p>0.05) |
| | pMAPK | Yes (p<0.05)<br>*Greater in females in aged-EVs group* | No (p>0.05) |
| | AP-1 | No (p>0.05) | No (p>0.05) |
| | TNF-α | Yes (p<0.05)<br>*However, post-hoc tests did not differ between males and females for all groups* | No (p>0.05) |
| | IL-8 | No (p>0.05) | No (p>0.05) |
| ***Mediators and end products of cGAS-STING activation*** | Total cGAS | No (p>0.05) | No (p>0.05) |
| | p-STING | No (p>0.05) | No (p>0.05) |
| | p-TBK1 | No (p>0.05) | No (p>0.05) |
| | p-IRF3 | Yes (p<0.05)<br>*Greater in females in aged-EVs group* | No (p>0.05) |
| | IFN-α | No (p>0.05) | Yes (p<0.05)<br>*Both males and females showed positive response* |
| **Mediators of JAK-STAT signaling activation** | p-JAK1 | No (p>0.05) | No (p>0.05) |
| | p-JAK2 | No (p>0.05) | No (p>0.05) |
| | p-STAT1 | No (p>0.05) | No (p>0.05) |
| | p-STAT3 | No (p>0.05) | No (p>0.05) |
| **Interferon Stimulatory genes** | IRF1 | No (p>0.05) | No (p>0.05) |
| | ICAM1 | No (p>0.05) | No (p>0.05) |
| | CD47 | No (p>0.05) | No (p>0.05) |
| | IRF5 | No (p>0.05) | No (p>0.05) |
| | Alcam | No (p>0.05) | No (p>0.05) |
| | IRF7 | No (p>0.05) | No (p>0.05) |
| | IFNγ | No (p>0.05) | No (p>0.05) |
| | TNF | No (p>0.05) | No (p>0.05) |
| ***Novel object (NO) discrimination index*** | Novel Object Recognition Test | No (p>0.05) | No (p>0.05) |

| | | | |
|---|---|---|---|
| ***Object in the novel location (OINP) discrimination index*** | Object Location Test | No (p>0.05) | No (p>0.05) |

**2.12. Supplementary Table 3**

**List of Genes in Microglial Clusters 1 and 2**

| **Genes in Microglial Cluster 1** | | |
|---|---|---|
| *Cmss1* | *Gm19951* | *Tmtc2* |
| *Filip1l* | *mt-Nd6* | *Gm15564* |
| *Il31ra* | *Gm48099* | |
| *Taco1* | *D130009I18Rik* | |

| **Genes in Microglial Cluster 2** | | | |
|---|---|---|---|
| *Ivns1abp* | *Ttc17* | *4931422A03Rik* | *Tasp1* |
| *8030442B05Rik* | *Trmt1l* | *Nme7* | *Miga2* |
| *Capn3* | *Dcaf17* | *D330023K18Rik* | *Inafm2* |
| *Abl1* | *Spopl* | *Vangl2* | *Traf3ip1* |
| *Gpr155* | *Ralgps1* | *Il15ra* | *Ankrd16* |
| *Mlph* | *Golga1* | *Hacd1* | *Gm37233* |
| *Pard3b* | *Itpripl1* | *Dolk* | *Fastkd1* |
| *2610203C22Rik* | *Tspan18* | *Col6a3* | *Disp1* |
| *Ccdc93* | *Hnmt* | *Ubox5* | *Bivm* |
| *Garnl3* | *Kcnip3* | *Usp20* | *Carf* |
| *Gm4258* | *Acsl3* | *Ino80dos* | *Stradb* |
| *A830008E24Rik* | *Cstf3* | *Atp1a2* | *Or5m3b* |
| *Mbd5* | *Acvr2a* | *Nemp2* | *Ttc21b* |
| *Slc40a1* | *Gpatch2* | *Zer1* | *Dusp27* |
| *Ggta1* | *Pced1a* | *Cry2* | *Sccpdh* |
| *Rabgap1* | *Gm27003* | *Rbm45* | *Utp25* |
| *Vis1* | *Usp21* | *Harbi1* | |
| *Ccnt2* | *Cep152* | *Ikzf2* | |
| *Pdk1* | *Chst14* | *Elp4* | |
| *Acvr1* | *Ppp1r12b* | *Slamf1* | |
| *Nr6a1os* | *Slc39a12* | *Hibch* | |
| *Eng* | *Zbed6* | *Slc1a2* | |
| *Kansl1l* | *Dtwd1* | *AI597479* | |
| *Nos1ap* | *Madd* | *Qser1* | |
| *Ralgps2* | *Bbs5* | *Poglut2* | |
| *Dis3l2* | *Tsga10* | *Accs* | |
| *Vcpip1* | *Yod1* | *Pigm* | |
| *Ubr1* | *Osgepl1* | *A130010J15Rik* | |

### 2.13. Supplementary Table 4

**List of Genes in Microglial Cluster 3**

| Genes in Microglial Cluster 3 | | | |
|---|---|---|---|
| *Il1r2* | *Ctse* | *Zfand2b* | *Gpc1* |
| *Il18rap* | *Sp100* | *Gm57177* | *Cnnm4* |
| *Stat4* | *Tuba4a* | *Flvcr1* | *Nhej1* |
| *Mfsd6* | *Coq10b* | *Hjurp* | *Gm7694* |
| *Stk17b* | *Pam* | *Gm16587* | *2900060B14Rik* |
| *Cxcr2* | *Gm19705* | *B4galt3* | *A430105J06Rik* |
| *Slco4c1* | *Suco* | *Inpp1* | *Ptp4a1* |
| *Cxcr4* | *Ppp1r42* | *Gm7160* | *Sag* |
| *Cd55* | *Adipor1* | *Panct2* | *Hsd17b7* |
| *Chil1* | *Degs1* | *Ctdsp1* | *Mpzl1* |
| *Rgs18* | *C130026I21Rik* | *Cpa6* | *Capn2* |
| *Ptgs2* | *Hdac4* | *Mndal* | *9430060I03Rik* |
| *Ptgs2os2* | *Neurl3* | *Klhl12* | *Gm57176* |
| *Sell* | *Niban1* | *Ptpn7* | *Gm57109* |
| *F5* | *Gm41914* | *Gm33887* | *Ikbke* |
| *H3f3a* | *Inpp4a* | *Nab1* | *Mreg* |
| *Lbr* | *Atg9a* | *Gm16083* | |
| *1700047M11Rik* | *St8sia4* | *Dcaf6* | |
| *4930523C07Rik* | *Mapkapk2* | *Arid5a* | |
| *Pbx1* | *Il18r1* | *Olfml2b* | |
| *Ncf2* | *Qsox1* | *Rpe* | |
| *Prdx6* | *Lrrfip1* | *Ifi203* | |
| *Serpinb10* | *Rnasel* | *Per2* | |
| *Sp140* | *Agap1* | *Mpc2* | |
| *Nabp1* | *Fcgr4* | *Kdm5b* | |
| *Btg2* | *Sde2* | *Fn1* | |
| *Arpc5* | *Tbc1d8* | *Cd244a* | |
| *A530032D15Rik* | *Hlx* | *Gm15832* | |

### 2.14. Supplementary Table 5

### List of Genes in Microglial Cluster 4

| **Genes in Microglial Cluster 4** | | | |
|---|---|---|---|
| *Atf3* | *Bcl2* | *Chchd5* | *Rpp38* |
| *Rgs1* | *Ephx1* | *Niban2* | *2310009B15Rik* |
| *Ier5* | *Nrp2* | *Phpt1* | *Patl2* |
| *Il1a* | *Neurl3* | *Zfp770* | *Nsmf* |
| *Slc25a25* | *Dpp7* | *B930036N10Rik* | *Gm31728* |
| *Mapkapk2* | *Gm57148* | *Timm10* | *Raph1* |
| *Btg2* | *Arl5b* | *Ifi213* | *Mrc1* |
| *Rab7b* | *Pcna* | *Ralgds* | *Ckap2l* |
| *St8sia6* | *Igsf8* | *Kynu* | *Cd244a* |
| *Ifi204* | *Otud1* | *Gm57182* | *Nr4a2* |
| *Optn* | *Rxrg* | *Esrrg* | *Ikbke* |
| *Ifi207* | *Snhg6* | *D730003I15Rik* | *Tmem177* |
| *Tank* | *Ifi211* | *Stk11ip* | *2900060B14Rik* |
| *Pfkfb3* | *Tmbim1* | *Il1r1* | *Itga4* |
| *Pdcd1* | *Uap1l1* | *B3gnt7* | *Fn1* |
| *Ly9* | *Chst1* | *Cacna1s* | *Mndal* |
| *Etl4* | *Speg* | *Traf2* | |
| *Fmn1* | *Tmem163* | *Ifi206* | |
| *Coq10b* | *Prr5l* | *Catip* | |
| *Slamf7* | *Psen2* | *Kif1a* | |
| *Abl2* | *Ifi209* | *Fzd7* | |
| *Slc23a2* | *Nsun6* | *Dusp2* | |
| *Gm13391* | *Cfap45* | *Zdhhc12* | |
| *Tagln2* | *Pgap1* | *Gm14023* | |
| *Ass1* | *Nuak2* | *Dusp10* | |
| *Gm37168* | *Cdk18* | *Coq8a* | |
| *Slamf8* | *Pogk* | *Spata5l1* | |
| *Garin4* | *E030042O20Rik* | *Slc30a1* | |

### 2.135 Supplementary Table 6

### List of Genes in Microglial Cluster 5

| Genes in Microglial Cluster 5 | | | |
|---|---|---|---|
| *Sgo2a* | *Mfsd6* | *Atp2b4* | *Coq10b* |
| *Tuba4a* | *1810006J02Rik* | *Agap1* | *Rev1* |
| *Gpc1* | *Ptgs2* | *Arpc5* | *Snrpe* |
| *Ptpn7* | *Ptgs2os2* | *Gm16083* | *Nab1* |
| *Kif14* | *Panct2* | *Stk16* | *Sox13* |
| *Aspm* | *Ikbke* | *Ctdsp1* | *Gm57176* |
| *Slco4c1* | *Sp140* | *Klhl12* | *Gm29170* |
| *Serpinb10* | *Inpp1* | *Gm41914* | *Tmem81* |
| *2810408I11Rik* | *Ankmy1* | *B230216N24Rik* | *Nucks1* |
| *Ppp1r42* | *C130026I21Rik* | *Tbc1d8* | *Ndufb3* |
| *Ube2t* | *Per2* | *Ap1s3* | *Adhfe1* |
| *Chil1* | *Hjurp* | *Fn1* | *Gm29570* |
| *Rgs18* | *Rufy4* | *Gm10552* | *Gm15832* |
| *Chit1* | *Serpinb2* | *Mterf4* | *2900060B14Rik* |
| *Csrp1* | *Zfand2b* | *Gm57162* | *Clasp1* |
| *Stat4* | *Stk17b* | *Sned1* | *Utp14b* |
| *Bard1* | *Atg9a* | *Gm57188* | |
| *Pam* | *Nabp1* | *9430060I03Rik* | |
| *Il18rap* | *Tex30* | *Inpp4a* | |
| *Cxcr2* | *Mcm6* | *Hdac4* | |
| *Il1r2* | *Pask* | *Gm7160* | |
| *Gm57177* | *Nhej1* | *Trak2* | |
| *Cd55* | *Gm57169* | *Mreg* | |
| *Il18r1* | *Ctse* | *D630023F18Rik* | |
| *A530032D15Rik* | *Sp100* | *Asnsd1* | |
| *Cxcr4* | *Rpe* | *Uchl5* | |
| *Cpa6* | *Dtymk* | *Atg4b* | |
| *Gm19705* | *Mybl1* | *Gm57183* | |
